# WAH-*i*: Optimising Microphone Array Geometry for Customised Localisation Accuracy

**DOI:** 10.64898/2026.02.07.704547

**Authors:** Ravi Umadi

## Abstract

1. Accurate spatial localisation of free-flying echolocating bats is foundational for resolving fine-scale flight and echolocation behaviour, prey interception, and spatial decision-making in natural environments. Acoustic localisation using microphone arrays is widely employed for this purpose, yet array geometries in field studies are typically chosen heuristically rather than systematically optimised. As portable multichannel ultrasonic recording systems become increasingly accessible, principled design guidelines are needed to ensure reliable localisation performance under practical deployment requirements.
2. I introduce an iterative array optimisation algorithm that designs microphone geometries by maximising localisation reliability within a predefined three-dimensional field of interest. The method evaluates candidate geometries using simulated acoustic emissions and time-difference-of-arrival localisation, quantifying performance as a volumetric pass rate: the proportion of source locations that meet a user-defined accuracy threshold. Microphone positions are iteratively perturbed and accepted based on improvements to this task-level metric, while enforcing practical constraints on array aperture, inter-sensor spacing, and deployability.
3. Across canonical polyhedral geometries, random initialisations, and arrays comprising four to twelve microphones, optimisation consistently produced rapid early gains followed by geometry-dependent performance plateaus under the specified stopping criterion and iteration budget. Under fixed-aperture constraints, increasing the microphone count yielded diminishing returns, and optimised low-order arrays — particularly four-microphone configurations — matched or exceeded the volumetric localisation performance of higher-order arrays with suboptimal geometry. Analysis of optimisation trajectories further revealed that convergence dynamics scale with array order, whereas volumetric performance under the tested conditions is dominated by geometry rather than sensor number.
4. These results demonstrate that array geometry is the primary determinant of volumetric localisation reliability, and that efficient, portable arrays can be systematically designed using optimisation rather than heuristic rules. The proposed framework is broadly applicable to bioacoustic localisation problems beyond echolocating bats, including avian tracking, passive acoustic monitoring, and conservation-oriented sensing, and provides a general approach for designing task-optimised acoustic sensor arrays for a wide range of applications.

## 1 INTRODUCTION

Acoustic signals provide a non-invasive means of detecting animals and investigating their behaviour in environments where direct observation is difficult. Interpreting these signals often requires knowing where they originated, particularly when reconstructing movement, distinguishing simultaneously vocalising individuals, or relating vocal activity to habitat use. Across taxa, microphone arrays support acoustic localisation and source separation in behavioural ecology, biodiversity monitoring, and conservation (Blumstein et al., 2011; Khan et al., 2025; Okamoto & Oguma, 2025; Rhinehart et al., 2020; Tolkova & Klinck, 2022; Umadi, 2025b; Vallejo & Taylor, 2009; Verreycken et al., 2021). The reliability of the resulting biological inference therefore depends heavily on resolving its source position at the spatial scale relevant to the behaviour or monitoring objective.

Echolocating bats provide a particularly informative application because they navigate and forage in complex three-dimensional environments by emitting ultrasonic calls and analysing the returning echoes (Griffin, 1944, 1958; Griffin & Galambos, 1941; Schnitzler, 1968). This active sensing modality enables precise spatial perception without relying on vision and has made bats an important model system for studying acoustically guided spatial behaviour, sensory ecology, and predator–prey interactions in the wild (Stidsholt, 2025; Surlykke & Nachtigall, 2014). Quantitative investigation of these behaviours critically depends on the ability to localise animals accurately in three dimensions over ecologically relevant spatial scales (Gaudette & Simmons, 2014; Koblitz, 2018; Umadi, 2025c).

Many behavioural questions in bat ecology require centimetre-scale localisation accuracy across observation volumes spanning several cubic metres (Gaudette & Simmons, 2014; Umadi, 2025c). Examples include reconstructing flight trajectories during prey capture (Umadi & Firzlaff, 2025), estimating spatial decision rules during obstacle negotiation (Falk et al., 2014), and resolving fine-scale interactions between conspecifics or between bats and prey (Chiu et al., 2010). Accurate source localisation is also required to estimate emitted source levels and sonar-beam directionality after accounting for source–receiver geometry, geometric spreading, and frequency-dependent atmospheric attenuation (de Framond et al., 2023; Surlykke & Kalko, 2008; Surlykke et al., 2009). Achieving such precision under field conditions is challenging. Optical tracking systems are constrained by lighting requirements, line of sight, and restricted working volumes, while satellite-based positioning generally lacks the spatial and temporal resolution required for small, fast-moving animals. Acoustic localisation using microphone arrays therefore represents a particularly practical and widely used approach for tracking echolocating bats in natural environments.

Microphone-array systems estimate source position from the time differences of arrival (TDOA) of acoustic signals across spatially distributed sensors. This approach is well suited to nocturnal field studies, provides high temporal resolution when combined with high sampling rates, and can be tailored to species-specific call characteristics. In indoor and outdoor experiments, microphone arrays have been used to quantify sonar-beam directionality and active control of the bats’ sensory field of view (Brinkløv et al., 2011; Eitan et al., 2022; Kounitsky et al., 2015; Surlykke et al., 2009); measure emitted source levels and their adjustment during foraging and prey attacks (de Framond et al., 2023; Lewanzik & Goerlitz, 2018); reconstruct prey-interception trajectories and relate biosonar timing to sensorimotor transitions (Fujioka et al., 2016; Sumiya et al., 2017; Umadi & Firzlaff, 2025); examine how call production is coordinated with wingbeat, flight velocity, and prey distance (Jakobsen et al., 2025; Xia et al., 2025); and test how acoustic-flow cues regulate flight speed (Haron et al., 2026). Broader array-based approaches to studying bat echolocation in the laboratory and field have been reviewed by Gaudette and Simmons (2014) and Koblitz (2018). Despite these diverse applications, the geometric design of microphone arrays in biological research remains largely *ad hoc*, guided by heuristic symmetry or practical convenience rather than systematic optimisation.

Field deployments impose additional constraints that complicate array design. The number of microphones is limited by cost, synchronisation complexity, power consumption, and data throughput, often restricting systems to fewer sensors. Physical constraints, such as mounting structures, portability, and microphone placement, further limit design choices. Environmental factors, including wind, temperature gradients, and multipath reflections, introduce additional sources of localisation error. These constraints create a fundamental trade-off: increasing array aperture improves geometric precision, but larger arrays are harder to deploy and maintain. This motivates a central practical question: given a fixed microphone budget and physical constraints, what array geometry maximises localisation reliability over a target observation volume?

From a mathematical perspective, TDOA-based localisation is inherently geometry-dependent. Relative arrival times are estimated from cross-correlations of microphone signals and converted into source position estimates via multilateration (Knapp & Carter, 1976; Smith & Abel, 1987; Spiesberger, 1999, 2001; Spiesberger & Fristrup, 1990). While timing precision depends on factors such as sampling rate and signal-to-noise ratio (Carter, 1987; Knapp & Carter, 1976; Spiesberger & Fristrup, 1990), the resulting position error is strongly modulated by sensor geometry (Chan & Ho, 1994; Smith & Abel, 1987). For a fixed timing uncertainty, the same array can yield highly accurate localisation in some regions of space but substantially larger or numerically unstable position estimates in others. This spatial heterogeneity is commonly described by geometric dilution of precision (GDOP), which captures how geometry amplifies timing errors as a function of source position.

A substantial body of work has addressed sensor placement using analytical measures of estimation uncertainty. Approaches based on the Cramér–Rao lower bound seek geometries that increase Fisher information and reduce the minimum achievable estimation variance (Bishop et al., 2010; Ho & Vicente, 2008; Isaacs et al., 2009). Related analyses use GDOP and other geometry-dependent error measures to characterise how sensor arrangement affects localisation precision (Chan & Ho, 1994; Smith & Abel, 1987; Spiesberger, 2001). These methods provide valuable theoretical insight, but generally rely on a specified noise model and local approximation of estimation uncertainty. They do not directly evaluate the fraction of a three-dimensional observation volume that satisfies an operational accuracy requirement after signal generation, delay estimation, and multilateration.

Alternative approaches exploit statistical performance models or convex relaxations. Direction-of-arrival studies have established analytical relationships among array geometry, covariance, and estimation uncertainty (Stoica & Nehorai, 1990), while convex methods can efficiently select sensors from a predefined set of candidate locations (Joshi & Boyd, 2009). Such relaxations are valuable when the design variables and constraints admit a tractable convex representation, but they do not directly address continuous microphone placement under the combined minimum-spacing, non-coplanarity, and thresholded volumetric-accuracy requirements considered here. Related placement methods optimise Fisher information for mobile range-based target tracking (Martínez & Bullo, 2006) or visual coverage over two-dimensional regions (Schwager et al., 2009); neither directly incorporates the timing estimation and volumetric error structure of acoustic multilateration.

Consequently, several gaps remain. First, many optimisation studies emphasise pointwise or averaged error measures rather than task-level measures of volumetric reliability. Second, little attention has been paid to geometry-dependent performance plateaus under task-specific constraints and finite computational budgets. Third, optimisation outcomes are rarely assessed across repeated runs, leaving it unclear whether the resulting solutions are robust or dependent on particular initialisations. Finally, few studies have evaluated array-optimisation strategies using signal and propagation conditions representative of bat echolocation.

Here, I address these gaps by framing microphone-array design as a constrained, task-oriented optimisation problem. Rather than minimising average localisation error, I evaluate performance using a *volumetric pass rate*: the fraction of source locations within a three-dimensional observation volume that can be localised within a predefined accuracy threshold. Array design is therefore formulated as an optimisation over microphone coordinates, subject to constraints on array extent, minimum inter-sensor spacing, and three-dimensionality. Although the aperture bound is straightforward, the pairwise minimum-spacing and non-coplanarity requirements produce a non-convex feasible region. The thresholded pass-rate objective is also non-smooth: small coordinate changes may leave the score unchanged or cause discrete changes as individual test locations cross the acceptance threshold. Geometry-dependent error amplification, near-field ambiguities, and localisation failures further produce multiple locally favourable regions (Chan & Ho, 1994; Smith & Abel, 1987; Spiesberger, 2001; Umadi, 2025c). These properties preclude a direct analytical solution and make objective gradients unavailable or unreliable. I therefore use derivative-free search, in which candidate microphone movements are evaluated through their resulting localisation performance without requiring derivatives of the objective.

To search this landscape, I extend the Array Widefield Acoustics Heuristic (Array WAH) (Umadi, 2025a, 2025c) to an inverse, optimisation-driven framework termed the Widefield Acoustics Heuristic–*inverse iterative* (WAH-*i*) (Umadi, 2026). WAH-*i* implements derivative-free co-ordinate search by iteratively perturbing microphone positions, evaluating each candidate through the forward localisation pipeline, and retaining movements that improve volumetric pass rate under the prescribed spatial and geometric constraints. Using signal parameters representative of aerial-hawking bats, I evaluated six canonical starting geometries (Fig. 1b) alongside randomly initialised arrays. Repeated runs were used to characterise convergence, robustness, and geometry drift in the presence of stochastic initialisation and stall-escape perturbations.

**Figure 1:**
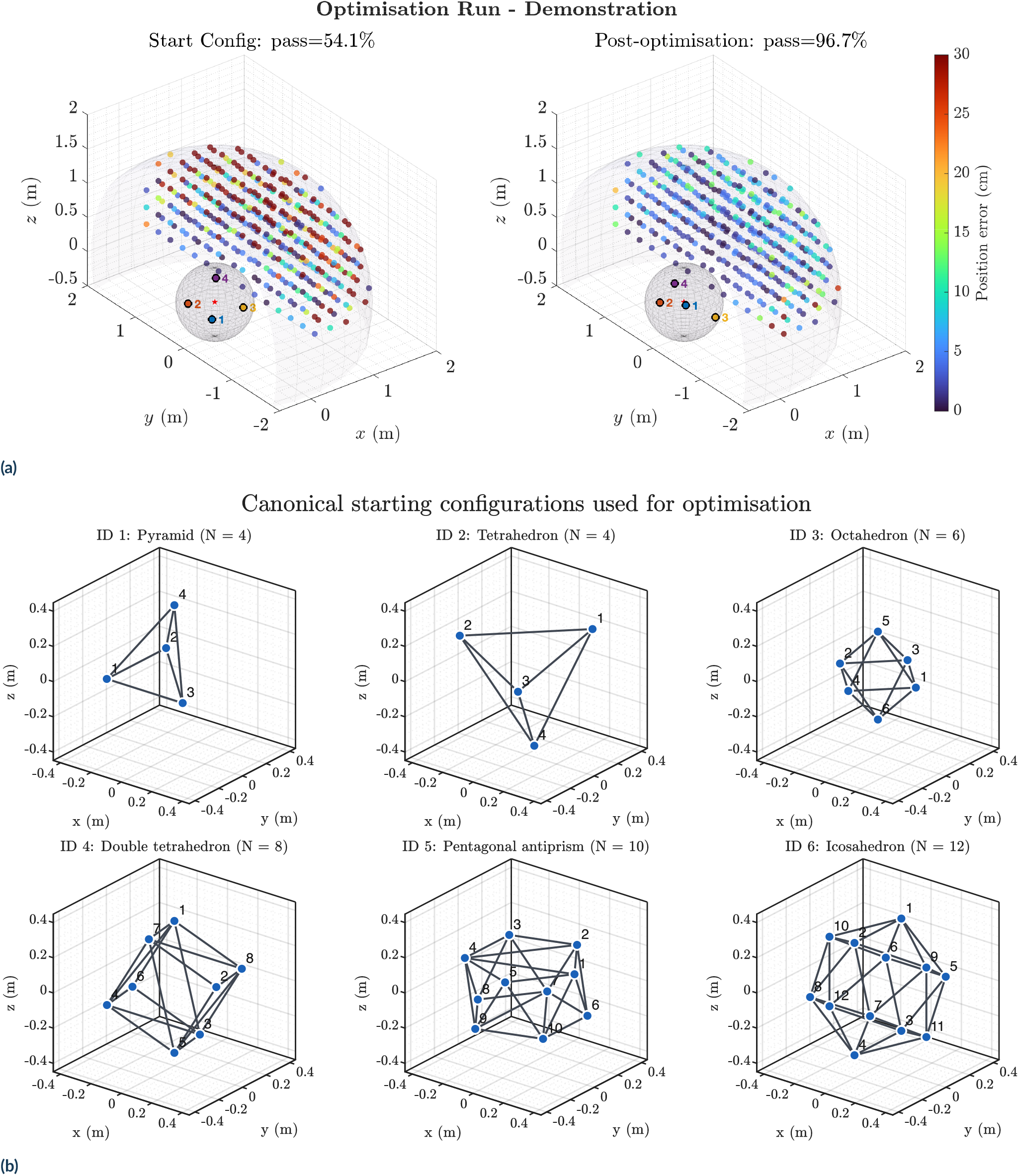
Representative optimisation outcome and starting configurations. **(a)** Localisation-error maps before and after optimisation for a representative run starting from the pyramidal configuration (configuration ID 1) illustrate how optimisation changes the spatial distribution of error. **(b)** The six canonical starting geometries are shown with their configuration IDs and microphone counts: (1) pyramid, (2) tetrahedron, (3) octahedron, (4) double tetrahedron, (5) pentagonal antiprism, and (6) icosahedron. Blue points indicate microphone positions, numbered by microphone index, and connecting lines show the edges of the corresponding convex hull. Coordinates are given in metres.

The results demonstrate that microphone array performance is governed less by sensor count *per se* than by how effectively a given geometry exploits the constraints of the task and sensing volume. By framing array optimisation in terms of volumetric reliability, economy, and responsiveness, this study provides a principled alternative to heuristic or scale-driven design choices that have traditionally guided bioacoustic localisation systems. The finding that carefully optimised low-order arrays can match or exceed the usable performance of denser configurations has immediate practical implications for field deployments, where portability, mechanical stability, power consumption, and ease of replication are often limiting factors.

More broadly, WAH-*i* treats array geometry as a design variable rather than a configuration inherited from convention. This principle extends to active and passive acoustic-sensing applications in which spatial accuracy, coverage, robustness, and system complexity must be balanced against explicit measurement requirements.

## 2 METHODS

I combined simulation-based localisation with iterative geometry optimisation to evaluate and refine microphone arrays for a specified three-dimensional field of accuracy under spatial and mechanical constraints. The following sections describe the performance criterion, signal and localisation model, optimisation procedure, experimental design, and analysis of the resulting configurations.

### 2.1 Optimisation formulation: task-level array design for volumetric TDOA localisation

I considered passive three-dimensional acoustic localisation of echolocation calls using time-difference-of-arrival (TDOA) measurements from a compact microphone array. Let an array configuration be represented by a matrix M ∈ ℝ^*N* ×3^, whose rows m_*i*_ = [*x*_i_, y_*i*_, *z*_*i*_] ^T^ give the Cartesian co-ordinates of the *i*-th microphone. Array performance was evaluated over a discrete set of candidate source positions *G* ⊂ℝ^3^ sampling the target observation volume. For a source at s ∈ *G*, localisation yields an estimate 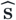, and I define the Euclidean position error

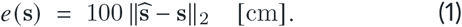

Because field tracking often hinges on whether localisation meets an operational accuracy requirement rather than on mean error alone, I used a task-level *pass–fail* criterion: a source location was deemed a *pass* if

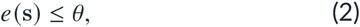

with acceptance threshold *θ* = 15 cm. This value was selected as a representative task-level requirement for comparing geometries within the specified observation volume, rather than as a universal threshold for bioacoustic localisation. In WAH-*i, θ* is user-defined and should be chosen according to the spatial scale of the biological question. A smaller value imposes a more stringent optimisation objective but does not itself increase the cost of evaluating a candidate geometry; instead, it can increase the number of iterations required to attain the desired volumetric pass rate. The primary performance metric was the *volumetric pass rate*, defined as the fraction of valid evaluation points meeting the criterion:

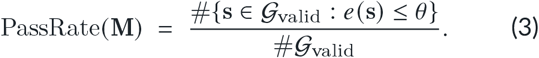

Here *G*_valid_ denotes the subset of evaluation points retained after considering the exclusion conditions (e.g. source positions too close to microphones; see below). In addition to positional error, I computed an angular error relative to a reference microphone by comparing estimated and true azimuth–elevation directions and reporting the Euclidean difference in the (az, el) plane (degrees). Position error was used for optimisation and for the pass criterion, while angular error served as a complementary performance descriptor (analysed in Fig. 5 and Fig. 7).

Array design was posed as a constrained nonlinear optimisation problem: find microphone coordinates M that maximise localisation reliability over *G* subject to physical boundary constraints. Specifically, all candidate geometries were required to satisfy (i) a bounded aperture constraint,

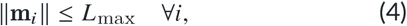

with ***L***_max_ = 0.50 m; and (ii) a minimum inter-microphone spacing constraint,

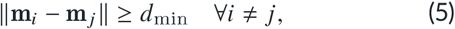

with *d*_min_ = 6.25 cm. This minimum separation was imposed as a practical construction constraint: it reserves space for microphones, mounting components, and supporting arms, reduces the likelihood of physical occlusion, and prevents the optimiser from exploiting near-coincident microphone positions that would be difficult to reproduce experimentally. The value is user-adjustable and should be adapted to the dimensions of the intended hardware. Candidate geometries were also required to be non-coplanar (numerically rank-3 after centring), ensuring a fully three-dimensional array capable of resolving elevation.

Although these constraints define a bounded set of admissible array geometries, the resulting optimisation problem is strongly non-convex. Small changes in microphone placement can produce abrupt changes in localisation performance due to GDOP, near-field effects, and occasional failures in time-difference-of-arrival estimation and nonlinear multilateration. In addition, performance was evaluated using a discrete pass–fail criterion rather than a smooth cost function, further fragmenting the objective landscape. As a result, the optimisation surface contains multiple local optima and discontinuities, making gradient-based or convex methods unsuitable and motivating the use of direct, derivative-free search strategies.

### 2.2 Evaluation grid and spatial sampling of the observation volume

Performance was evaluated on a three-dimensional lattice of candidate source positions with step size 0.25 m, spanning a shell-like region around the array centre. The outer radius was *R* = 2.0 m and an inner exclusion radius *R*_in_ = 0.30 m prevented evaluation arbitrarily close to the array. The inner exclusion is important because both acoustic time-delay estimation and multilateration can become ill-conditioned in extreme near-field scenarios (e.g. when the source is close to one sensor). In addition, I restricted evaluation to a forward, upper sector of space to reflect typical field arrangements in which the array is placed below or beside the flight volume; thus, not all spatial octants contributed equally to the objective. Grid spacing determines the spatial resolution of the objective and is the principal computational scaling parameter in the present framework. For a fixed three-dimensional observation volume, the number of evaluation points scales approximately with the inverse cube of grid spacing; halving the spacing therefore increases the number of evaluated locations by approximately eightfold. Because this full-grid evaluation is repeated for every candidate microphone displacement, the 0.25 m spacing represents a compromise between resolving broad spatial differences among geometries and maintaining tractable batch computation.

For each evaluation point, localisation was performed only if the source-to-nearest-microphone distance exceeded the minimum safety distance (0.08 m). Points failing this criterion were excluded from summary statistics and pass-rate calculation. Numerical failures (e.g. invalid delays or solver non-convergence) at otherwise valid points were assigned a large fixed penalty error; this prevents such cases from being silently ignored while ensuring that failures reduce the pass rate.

### 2.3 Forward acoustics, TDOA estimation, and multilateration

Following the synthetic signal-generation and propagation procedure developed for Array WAH (Umadi, 2025c), I simulated frequency-modulated echolocation calls with parameters representative of insectivorous bats. For the present analyses, each signal comprised a 70–30 kHz FM downsweep sampled at *f*_*s*_ = 192 kHz, with a duration of 5 ms, and additive white Gaussian noise scaled to produce an effective signal-to-noise ratio of 40 dB. For a source at s, each microphone signal was generated by propagating the call with a fixed speed of sound *c* = 343 m s^−1^ and applying the corresponding time-of-flight delay. TDOAs were estimated relative to a designated reference microphone using normalised cross-correlation, with the delay chosen as the lag that maximised the correlation magnitude. These estimated delays define a set of range differences of the form

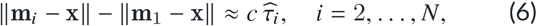

where x is the unknown source position and *τ*_i_ is the estimated TDOA relative to microphone 1. The source estimate s was obtained by solving the resulting nonlinear least-squares problem (hyperbolic multilateration). This procedure yields a localisation error vector over, from which pass rate and distributional summaries are derived. The implementation was built on the Widefield Acoustics Heuristic toolkit (Umadi, 2025a) (See Umadi (2025c) for the description of the original methodology).

### 2.4 Objective function for geometry optimisation

Optimisation targeted volumetric reliability while discouraging solutions that achieve a pass rate by tolerating large errors at the remaining points. The objective function was formulated as a goal-oriented scalarisation of competing criteria. Following the standard approaches in multi-objective optimisation (Boyd & Vandenberghe, 2004/ 2009; Miettinen, 1998), pass rate and mean localisation error were combined into a single objective using a linear weighting. To prevent domination by extreme outliers, the mean error term was normalised by a characteristic failure scale and capped, consistent with robust loss formulations used in inverse problems (Hampel, 1986; Huber, 1981). This construction reflects the operational goal of maximising volumetric reliability while weakly discouraging solutions that achieve high pass rates by tolerating extreme errors at a minority of points. I therefore maximised a scalar objective combining pass rate with a weak penalty on mean positional error,

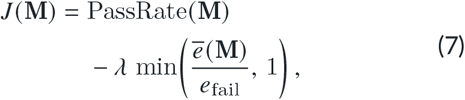

where *ē* (M) is the mean error over valid points, *e*_fail_ is the fixed penalty assigned to numerical failures, and *λ* is a small weighting factor. This form prioritises increasing the fraction of the working volume meeting the operational threshold while retaining sensitivity to broad error inflation and failure modes.

The choice of *λ* governs the trade-off between *coverage* and *accuracy margin*. For small values of *λ*, optimisation is dominated by pass-rate maximisation, favouring geometries that bring as many spatial locations as possible below the acceptance threshold, even if the resulting error distribution remains sharply peaked near that threshold. Increasing *λ* places greater emphasis on reducing the mean error among passing locations, encouraging solutions with improved robustness and larger safety margins at the expense of marginal reductions in pass rate. Importantly, because the penalty term is normalised and clipped, *λ* primarily influences behaviour near the pass–fail boundary rather than the treatment of extreme outliers.

The effectiveness of this trade-off is coupled to the spatial resolution of the coordinate-search procedure. Microphone positions are updated using a bounded coordinate descent with a linearly decaying step size,

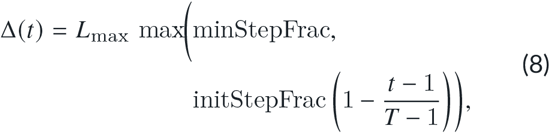

where *T* is the total number of optimisation iterations. Early iterations permit large exploratory movements that allow the optimiser to escape poor initial configurations, whereas later iterations perform fine-scale adjustments to refine geometry within a locally optimal basin.

Larger values of *λ* therefore benefit from sufficiently small terminal step sizes, as improvements in mean error typically require precise, local rearrangements of microphone positions rather than coarse geometric changes. Conversely, overly small initial step sizes can prevent the optimiser from escaping suboptimal basins regardless of penalty weighting. Together, *λ* and the step-size schedule define a coupled control mechanism that determines whether optimisation prioritises volumetric coverage, accuracy margin, or a balanced compromise between the two (also see Appendix B in Supplementary Material)

### 2.5 The WAH-*i* coordinate-search algorithm

The core coordinate updates use a deterministic direct-search strategy, augmented by stochastic stall-escape perturbations. The WAH-*i* algorithm performs iterative coordinate refinement by proposing candidate displacements for individual microphones, evaluating each candidate on the full grid *G*, and accepting the candidate that yields the greatest improvement in the objective *J*. This procedure is repeated across microphones in a cyclic schedule, ensuring that all sensors are revisited as the objective landscape evolves. A target pass rate of 95% was used as a task-sufficiency stopping criterion: optimisation terminated once at least 95% of valid grid locations met the 15 cm positional-error requirement, or when the maximum iteration budget was exhausted. This target limited computation after a configuration had achieved high volumetric reliability and should not be interpreted as an estimate of the maximum performance attainable by that geometry. Users seeking near-complete coverage may select a higher target, although increasingly stringent targets generally require longer runs, finer terminal step sizes, and potentially broader exploration of the optimisation landscape.

Candidate moves are generated along a fixed set of probe directions that include Cartesian axes and diagonals, as well as radial inward/outward directions relative to the array centre. Step size is governed by a scheduled decay: early iterations explore the space using larger steps to quickly exploit obvious improvements, whereas later iterations use smaller steps to refine local structure (see above). Candidate geometries are re-centred after each move to maintain a consistent array centre, and any microphone exceeding the aperture bound is projected back onto (or just within) the allowed maximum radius. Candidate moves violating the minimum spacing or non-coplanarity constraints are rejected without evaluation.

To avoid stagnation in shallow local optima, WAH-*i* includes a stall-detection mechanism. If successive iterations fail to improve the objective beyond a small tolerance, a stochastic perturbation (*shake*) is applied to the full geometry to promote escape from local basins. The perturbed geometry is subsequently repaired by projection and boundary checks; if the physical conditions cannot be restored, a new non-coplanar geometry is drawn. Under stringent accuracy or coverage requirements, improvements may become sparse as the search approaches a local plateau. Although stochastic perturbation permits exploration beyond such plateaus, it does not guarantee escape to a superior basin. Identifying more accurate configurations may therefore require larger iteration budgets, more independent runs, finer terminal step sizes, or more global search strategies.

### 2.6 Experimental conditions and repeated-run design

I evaluated the optimisation behaviour across the canonical three-dimensional starting geometries defined in Fig. 1b, spanning microphone counts *N*_mic_ = 4 to *N*_mic_ = 12, alongside randomly initialised arrays at matched *N*_mic_. The canonical configurations provide topology-informed geometric baselines, whereas random starts provide a practical reference for performance achievable without topology-informed initialisation.

Because direct-search optimisation in a non-convex landscape can be sensitive to initialisation and stochastic escape events, each configuration was optimised using five independent runs. This repeated-run design enables separation of systematic configuration-dependent performance differences from run-to-run optimisation variability, and supports statistical evaluation of convergence behaviour (Fig. 2b and Fig. 4).

**Figure 2:**
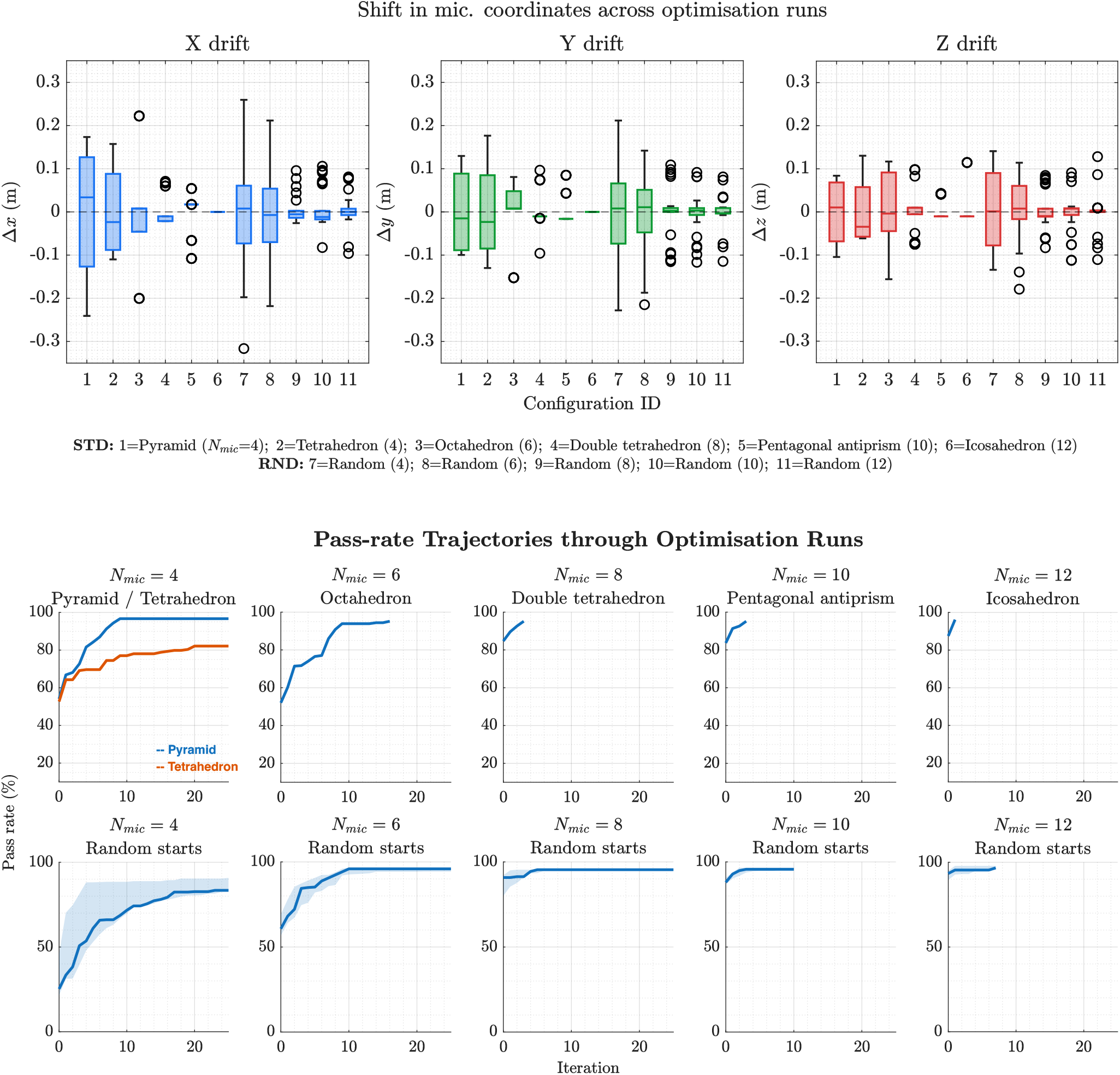
Optimisation dynamics and geometry changes. Microphone-array geometry was optimised by iteratively updating microphone coordinates to maximise the volumetric *pass rate*, defined as the percentage of test locations satisfying the predefined positional-accuracy threshold of `15` cm. **(a)** Distributions of the per-microphone coordinate changes (Δ*x*, Δy, and Δ*z*) between the initial and final configurations across five replicate runs, summarised by configuration ID. **(b)** Pass rate across optimisation iterations for canonical starting geometries (top row) and random starts (bottom row), grouped by microphone count (*N*_mic_). Solid lines show the median trajectory across five runs and shaded bands show the interquartile range. In the *N*_mic_ = `4` canonical panel, blue denotes the pyramid and orange denotes the tetrahedron; this colour convention is used consistently throughout the study.

**Figure 3:**
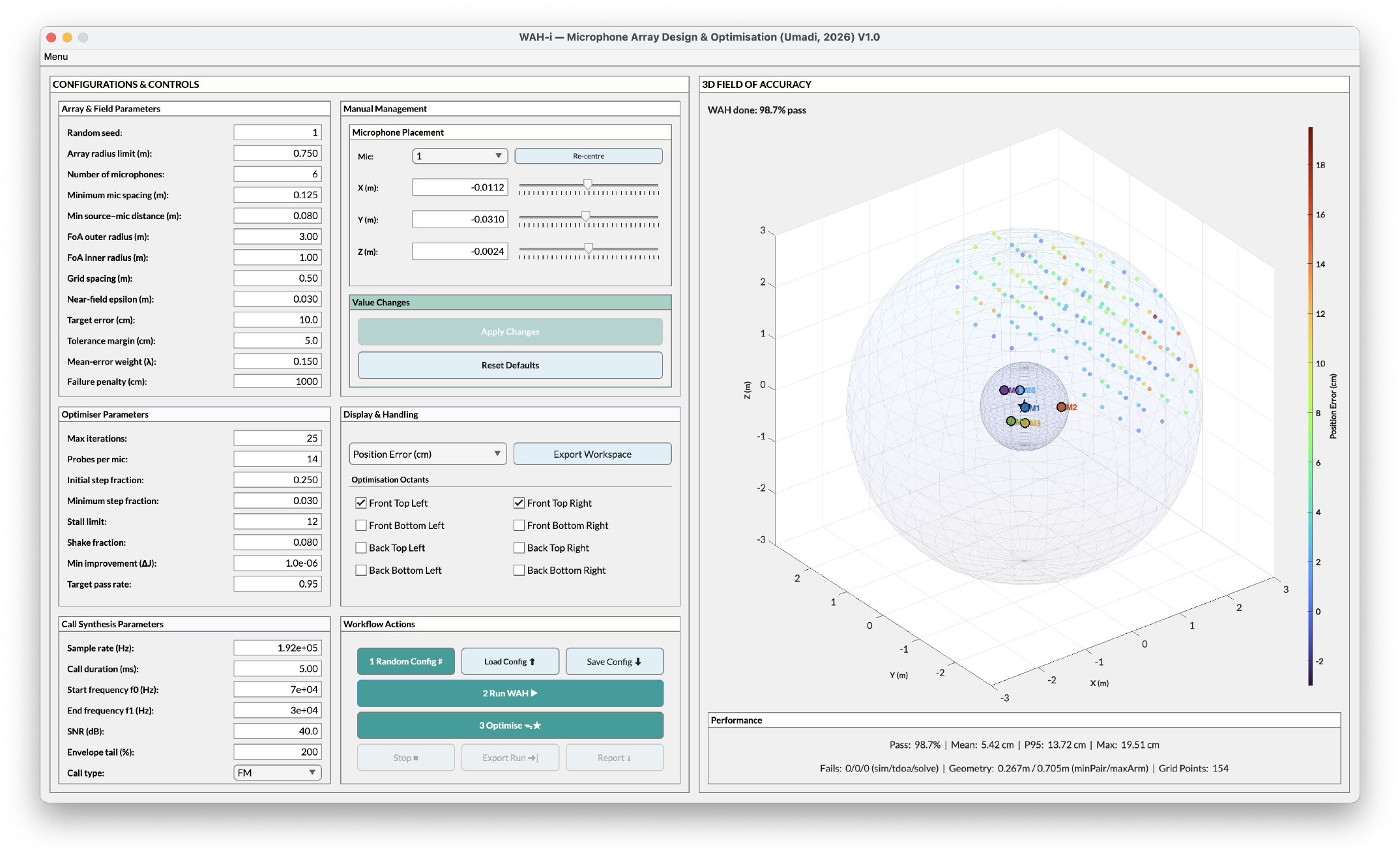
Interactive interface for array optimisation and performance exploration (WAH-*i* V1.0). The application provides an accessible entry point for exploring microphone array configurations under user-defined accuracy and spatial constraints. Users can specify the number of microphones, select or initialise array geometries, define aperture limits and localisation accuracy thresholds, and run the optimisation interactively. The interface visualises optimisation progress, the array geometry through optimisation, and the spatial distribution of localisation error within the evaluation volume.

### 2.7 Analysis of optimisation dynamics and localisation performance

Optimisation outcomes were examined at complementary levels to distinguish convergence behaviour, geometric change, and spatial and distributional performance.

#### Trajectory-level convergence and saturation

For each configuration, pass-rate trajectories were summarised across repeats using the median and uncertainty bands. Intermediate perturbation steps introduced during stall escapes were excluded to ensure consistent alignment (Fig. 2b).

#### Runtime diagnostics and geometry drift

To distinguish attainable performance from convergence speed, I compared the maximum and final pass rates and the iteration at which 95% of the run-wise maximum was first reached (Fig. 4). Per-microphone displacement vectors between the initial and final states quantified the magnitude and direction of geometry changes and whether improvement required substantial rearrangement (Fig. 2a).

**Figure 4:**
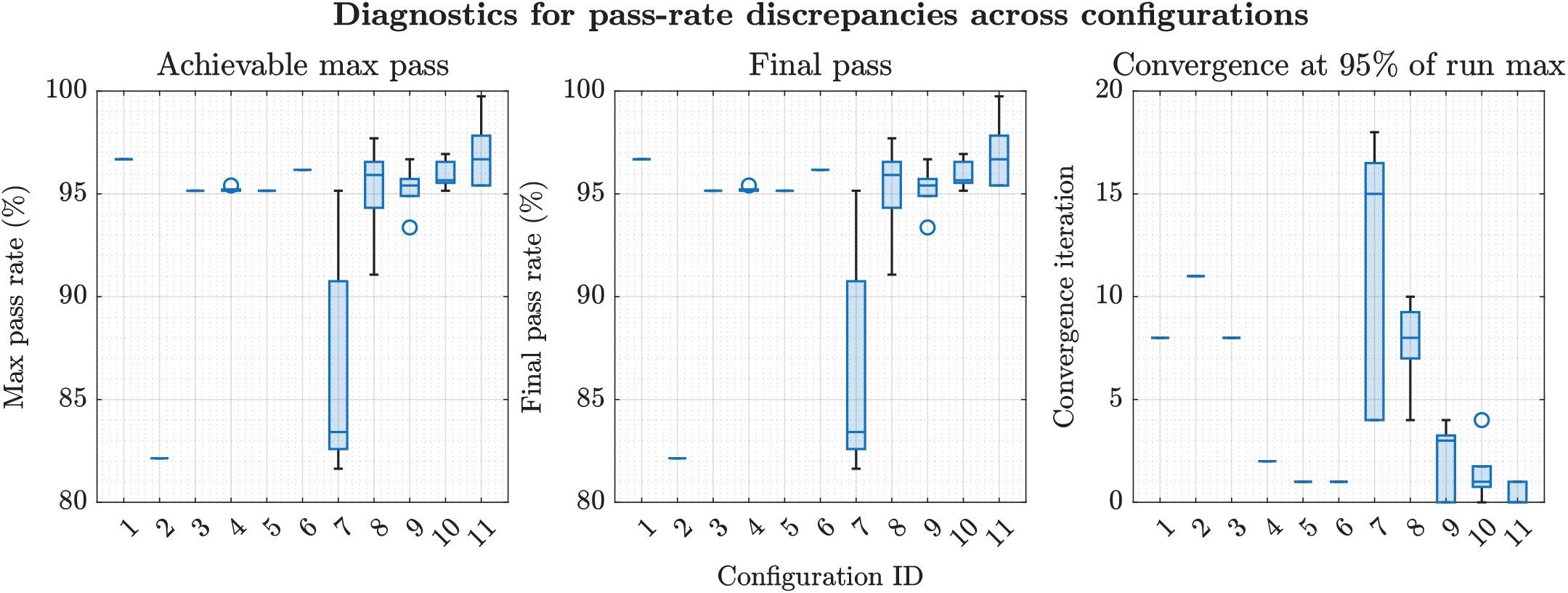
Runtime diagnostics underlying pass-rate discrepancies across microphone configurations. (Left) Distribution of the *achievable maximum pass rate* attained during optimisation, summarised across independent runs for each configuration. (Centre) *Final pass rate* at the end of optimisation, highlighting cases where convergence terminates below the run-wise maximum. (Right) *Convergence iteration*, defined as the iteration at which 95% of the run-wise maximum pass rate is first reached.

#### Spatial structure: error versus distance

Grid points were binned by radial distance from the array centre, and median error and 10–90% bands were calculated to quantify where distance-dependent degradation emerged under the fixed aperture constraint (Fig. 5).

**Figure 5:**
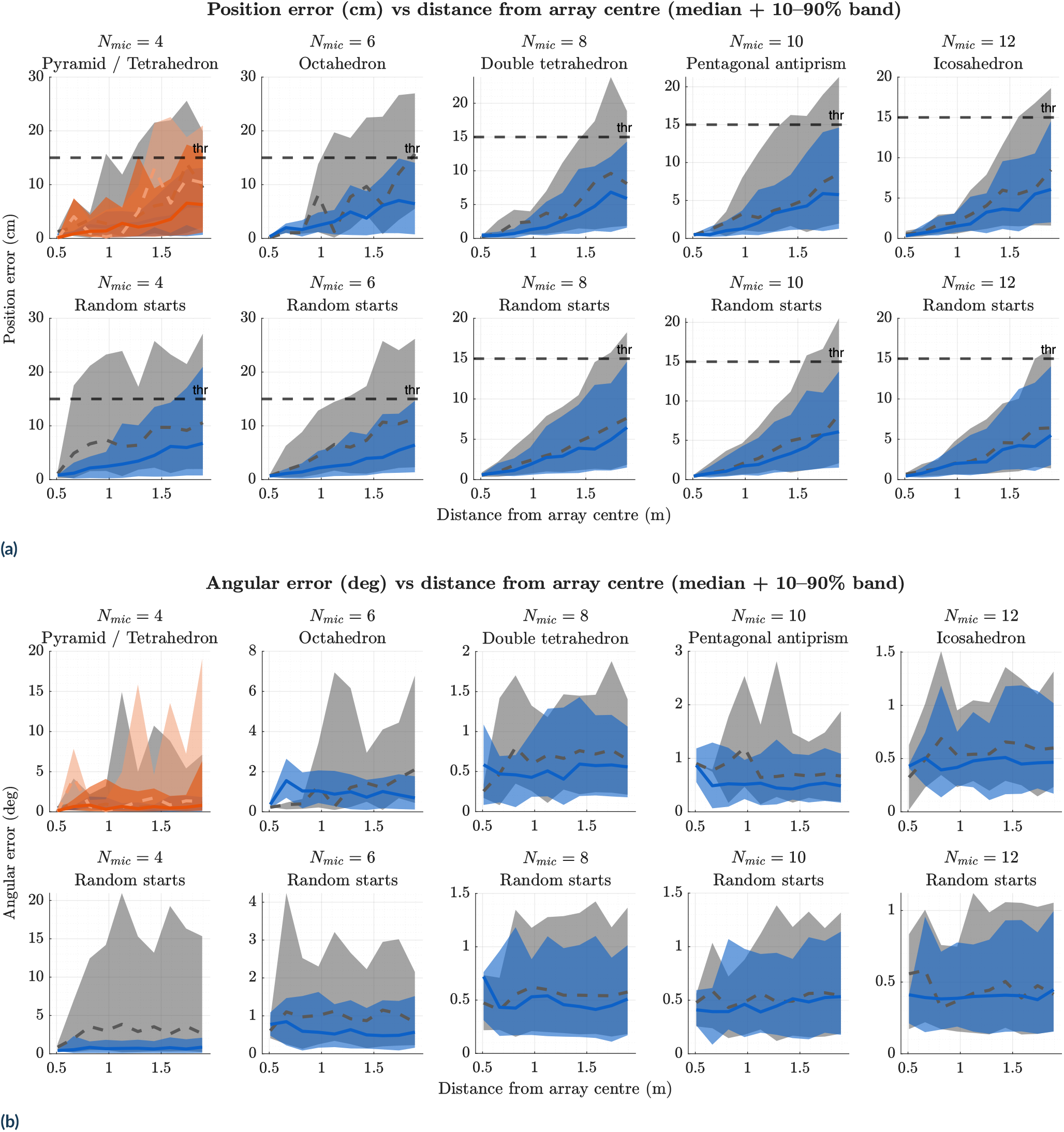
Optimisation improves spatial accuracy across distance. Localisation error is shown as a function of distance from the array centre for canonical starting geometries (top panels) and random starts (bottom panels), grouped by microphone count (*N*_mic_ = 4–12). Within each run, errors were grouped into radial-distance bins and summarised by the median and 10th–90th percentiles; the plotted curves and bands show the across-run medians of these bin-wise summaries. Grey dashed curves and lighter/grey shading represent performance before optimisation, whereas coloured solid curves and shading represent performance after optimisation. Vertical axis limits are adjusted by panel. **(a)** Positional error (cm) generally increases with distance, with geometry-dependent differences remaining most evident at larger radii; the optimisation objective and volumetric pass-rate calculation used a 15 cm positional threshold. **(b)** Angular error (degrees) exhibits a similar distance dependence but is less sensitive to array geometry.

#### Distributional performance before vs. after optimization

Errors on the same valid grid points before and after optimisation were compared using average-run histograms and robust statistics, including the 95th percentile (*p*_95_), to distinguish changes in typical accuracy from changes in the upper tail of each distribution (Figs. 6 and 7).

**Figure 6:**
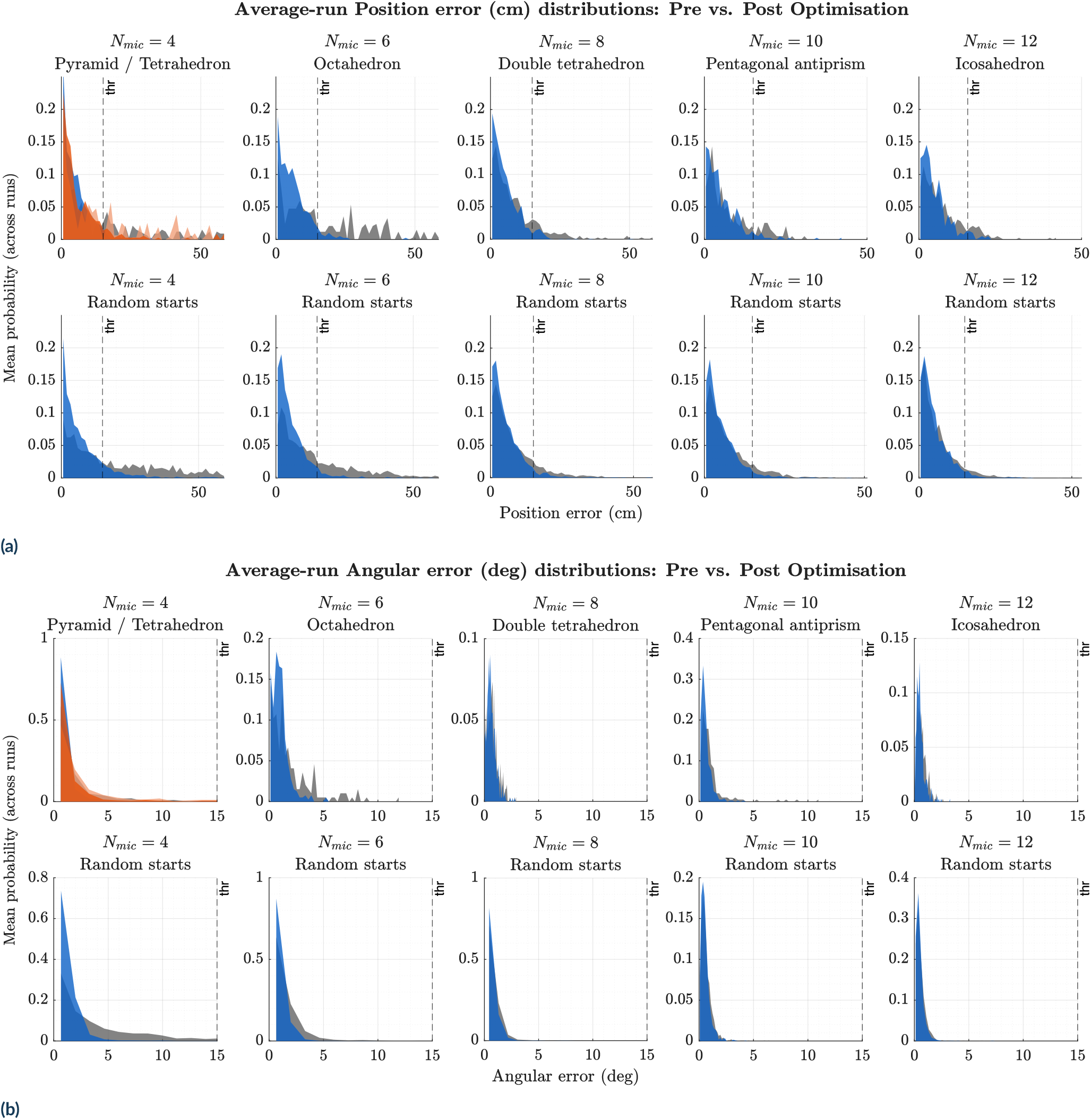
Localisation performance across array geometries. Average-run distributions of **(a)** positional error and **(b)** angular error are shown across canonical starting geometries and random initialisations. For each optimisation run, a probability-normalised histogram was calculated across valid source locations; the plotted distributions are the bin-wise means of these histograms across five replicate runs. Grey shading represents performance before optimisation and coloured shading represents performance after optimisation. Grey and blue denote the pyramidal configuration before and after optimisation, respectively, while light orange and orange denote the corresponding tetrahedral distributions in the four-microphone canonical panel. Vertical probability-axis limits are adjusted by panel. Values exceeding four times the displayed threshold were excluded from the histogram range to prevent extreme errors from obscuring the bulk of each distribution. The black vertical dashed lines indicate descriptive acceptance thresholds of 15 cm for positional error and 15^°^ for angular error; only the positional threshold was used in the optimisation objective and volumetric pass-rate calculation.

### 2.8 Software implementation and user-accessible optimisation tool

The optimisation and simulation pipeline was implemented in MATLAB R2023b (MathWorks, Natick, MA, USA) and is available through documented functions, scripts, and an interactive application (Fig. 3). Users can specify microphone count, initial geometry, aperture and spacing constraints, optimisation settings, and accuracy requirements without modifying the underlying code. Standalone executables generated with the MATLAB Compiler are provided for macOS and Windows (x64), allowing the same optimisation workflow to be used without a local MATLAB installation.

The complete versioned release, including the source code, analysis scripts, data, configuration files, and standalone application packages, is archived on Zenodo (Umadi, 2026). This combination supports reproducible batch analysis and exploratory use, while allowing researchers without programming experience to examine how geometry and task-specific constraints affect localisation performance before constructing or deploying a physical array.

## 3 RESULTS

Across all microphone counts and initialisation strategies, the optimisation procedure produced rapid early improvements in localisation performance, followed by clear saturation at configuration-specific pass-rate levels (Fig. 2). Pass-rate trajectories exhibited a consistent two-phase structure (Fig. 2b): an initial growth phase, in which performance increased steeply within the first 5–10 iterations, and a subsequent plateau phase, in which further coordinate updates yielded little or no additional improvement.

The early growth phase reflects the optimiser’s ability to efficiently exploit the most informative geometric degrees of freedom. In contrast, the plateau phase indicates the presence of structural limits beyond which further refinement cannot substantially increase the fraction of test locations meeting the accuracy criterion. Importantly, this saturation behaviour was highly reproducible across repeated runs and was observed for both canonical geometric initialisations and random starts, suggesting that convergence is governed by geometry-dependent feasibility rather than by optimisation instability or insufficient runtime.

Representative visualisations illustrate how optimisation reshapes spatial error structure, eliminating geometry-specific failure modes (Fig. 1a). Large contiguous regions of high error are reduced after optimisation. These starting residual structures differ systematically between geometries, but the optimisation almost entirely eliminates them.

The distribution of per-microphone coordinate changes further demonstrates that optimisation operates primarily through modest adjustments rather than radical rearrangements (Fig. 2a). Typical displacements were on the order of ± 0.1 m per axis, with no evidence for large-scale geometric reconfiguration. Together, these trajectory-level results show that optimisation is effective at rapidly improving performance, and that even denser arrays, although starting at a high pass rate, can benefit from configurational optimisation.

### 3.1 Geometry-dependent limits on achievable pass rate

Optimisation outcomes differed systematically across array geometries, with configurations reaching distinct performance plateaus under identical optimisation settings, stopping criteria, and repeated runs (Fig. 4). The distribution of run-wise maximum pass rates (Fig. 4, left) reveals clear performance stratification by array topology within the tested search budget. Compact and highly symmetric configurations, the tetrahedral array, consistently plateaued at substantially lower pass rates than more spatially distributed geometries, including pyramidal, antiprismatic, and higher-order polyhedral layouts.

Final pass rates closely tracked the corresponding run-wise maxima for most configurations (Fig. 4, centre), indicating that the optimiser generally converged reliably once a favourable region of the solution space was reached. Configurations exhibiting broader inter-run variability showed larger discrepancies between maximum and final performance, suggesting sensitivity to local optima rather than premature termination of the optimisation.

Convergence behaviour exhibited a clear dependence on configuration, but this dependence did not mirror the performance attained within the search budget. The iteration at which 95% of the run-wise maximum pass rate was first reached (Fig. 4, right) decreased systematically across configurations, with higher-order and random initialisations typically converging in fewer iterations than lower-order, highly constrained geometries. Notably, some of the fastest-converging configurations nonetheless achieved similar maximum pass rates, whereas several high-performing geometries required more iterations to approach their observed plateaus. This dissociation indicates that rapid convergence reflects a more tightly constrained optimisation landscape rather than superior localisation capability. Pass-rate differences therefore reflect configuration-dependent performance under the present constraints and stopping rules, rather than termination behaviour alone.

Across pooled runs, moderate correlations were observed between maximum achievable pass rate and optimisation dynamics. The maximum pass rate correlated positively with the mean optimisation step size (*r* = 0.64), and negatively with overall root-mean-square microphone displacement (*r* = − 0.53). These relationships indicate that higher-performing configurations tend to improve through coherent, moderately sized adjustments rather than through large cumulative geometric rearrangements. Importantly, extensive geometric drift was not predictive of superior performance. Together, these findings confirm that optimisation cannot compensate for topologically disadvantaged array designs; geometry selection therefore sets the fundamental limits on achievable localisation reliability.

### 3.2 Spatial structure of localisation accuracy

Optimisation improved localisation accuracy across most of the workspace, but systematic spatial structure remained evident after convergence (Fig. 5). For all configurations, positional error increased with radial distance from the array centre, reflecting persistent geometric dilution of precision. Optimisation primarily reduced errors within a geometry-dependent sensing volume rather than eliminating distance-dependent degradation.

Positional accuracy showed strong dependence on array geometry (Fig. 5a). Positional errors for compact and low-order configurations extended beyond the acceptance threshold near the edge (∼ 1.9 m), although the median remained within the threshold, whereas more spatially distributed arrays maintained acceptable accuracy. These differences define geometry-specific effective sensing radii that optimisation alone cannot exceed, constrained by array topology and aperture — a design consideration important for optimising far-field accuracy.

Angular error exhibited a similar radial trend but weaker sensitivity to geometry (Fig. 5b). However, angular accuracy generally improved and remained stable (median <1 deg) as the microphone count increased. This indicates that positional accuracy is more strongly constrained by array geometry under the present optimisation objective. Across all configurations, the 10–90% quantile bands for positional accuracy widened substantially at larger radii, revealing pronounced spatial heterogeneity in localisation reliability. Even after optimisation, peripheral regions of the workspace contained localised pockets of high error that could not be eliminated solely through coordinate refinement. However, the pass rate threshold was limited to 95; therefore, a more robust optimisation outcome can not be excluded. These results demonstrate that array geometry determines not only average accuracy but also the spatial extent over which reliable localisation can be achieved, and that these values can be optimised.

### 3.3 Optimisation reshapes error distributions

Baseline localisation performance varied widely across array geometries prior to optimisation, with large differences not only in pass rate but also in the structure of error distributions (Fig. 6). In several configurations—particularly random initialisations and low-order canonical geometries—baseline localisation was characterised by heavy-tailed error distributions, with failures dominating the upper percentiles (Supplementary Tables A1 and A2).

Optimisation consistently reshaped these distributions by reducing both median and upper-tail errors, although the relative magnitude of improvement differed among configurations (Fig. 7). Median positional error decreased substantially for the four-microphone canonical configurations, from 11.6 cm to 3.8 cm for the pyramid and from 13.0 cm to 4.2 cm for the tetrahedron, corresponding to reductions of approximately 67–68%. Improvements in median error were smaller for higher-order polyhedral arrays, decreasing from approximately 5–6 cm to 3–4 cm. Similar configuration-dependent shifts were observed for random initialisations once degenerate configurations were resolved.

**Figure 7:**
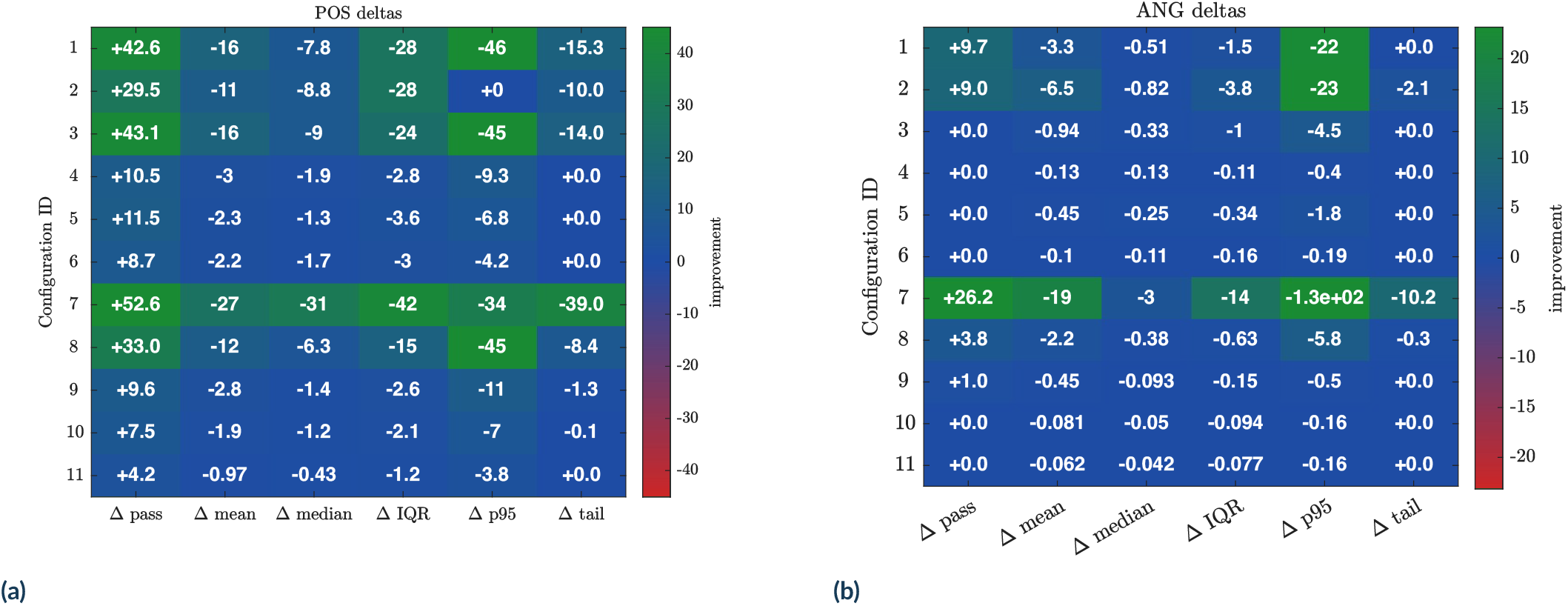
Optimisation-induced changes in localisation error. For each optimisation run, errors were evaluated on the same test grid at the first recorded iteration (*it* = 0; *before*) and at the selected *after* state (final iteration in the current analysis). Per-grid-point differences were computed as Δ*e* = *e*_after_ − *e*_before_ using the positional error (a) and the angular error (b). The differences were summarised using robust statistics, including the 95th percentile error (*p*_95_) to characterise tail behaviour. Distributions of Δ*e* are shown by configuration.

The most pronounced effects of optimisation nevertheless occurred in the upper tails of the error distributions.

In contrast, reductions in extreme positional error were substantial. The 95th percentile position error ∼ decreased by an order of magnitude in several cases, falling from values exceeding 1 m in poorly conditioned baseline geometries to ∼ 14–27 cm after optimisation. The most dramatic improvements occurred for random four-microphone arrays, where the 95th percentile positional error was reduced from approximately 5.9 m to 26 cm. Even for structured geometries with moderate baseline performance, such as the tetrahedron, the 95th percentile error dropped from 118 cm to 104 cm (Supplementary Table A1), while higher-order configurations converged to consistently low upper-tail errors near the imposed threshold. These large reductions in tail error account for the majority of the observed gains in volumetric pass rate.

Angular error exhibited a qualitatively similar but weaker response to optimisation. Median angular error decreased modestly across most configurations, typically from ∼ 0.6–1.5^°^ to ∼ 0.4–0.6^°^. Upper-tail angular errors were reduced substantially in configurations with poor baseline geometry—most notably random initialisations—where the 95th percentile angular error fell from tens or hundreds of degrees to values below 3^°^. However, for several canonical geometries, angular error distributions changed little after optimisation, and in some cases interquartile angular error increased slightly despite improved positional pass rates. Notably, for all configurations, the optimised medians fell below an average of 0.5 degrees (Supplementary Table A2).

This asymmetric response reflects the structure of the optimisation objective, which prioritises meeting a positional accuracy threshold with only a weak penalty on mean error. Optimisation therefore reallocates sensing precision toward eliminating severe positional failures near the pass boundary, rather than uniformly improving accuracy across the entire workspace. As a result, volumetric reliability improves substantially even when median errors change only modestly.

Together, these results demonstrate that the principal effect of optimisation is the suppression of rare but severe localisation failures that dominate volumetric performance metrics. While this strategy is highly effective for maximising usable sensing volume, it also highlights a fundamental trade-off: objective functions tuned for threshold-based reliability may not optimise angular accuracy or spatial uniformity. Alternative formulations with carefully chosen parameters may therefore be required when applications demand consistent precision rather than robust pass-rate performance.

### 3.4 Microphone count effects and diminishing returns

Increasing microphone count improved achievable volumetric pass rates, but gains remained sublinear and exhibited clear diminishing returns under fixed-aperture constraints. While higher-*N* arrays tended to reach higher absolute pass rates, the incremental benefit per additional microphone decreased rapidly beyond low-order configurations, consistent with the economy analysis (Table 3).

To quantify whether localisation performance or optimisation behaviour scales systematically with microphone count, I evaluated a power-law model of the form

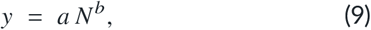

where *N* denotes the number of microphones and *y* is a run-wise summary metric. Fits were performed in log–log space using ordinary least squares, with uncertainty estimated via bootstrap resampling (2000 samples; 95% confidence intervals). The analysis was restricted to the random-start *N*-sweep (*N* = 4–12) to isolate the effect of microphone count from specific geometric symmetries.

As summarised in Table 1, both maximum and final volumetric pass rates exhibit a weak but positive scaling with microphone count (*b* ≈ 0.095), explaining approximately half of the variance in log space (*R*^2^ ≈ 0.51). While this indicates that higher-order arrays can, on average, achieve higher pass rates, the shallow exponent confirms that these gains increase slowly with *N* and fall far short of proportional scaling. In practical terms, doubling the number of microphones yields only marginal improvements in usable localisation volume.

In contrast, optimisation dynamics exhibit stronger, more systematic scaling with microphone count. The iteration at which the optimiser converges decreases steeply with *N* (*b* ≈ −2.10), indicating that higher-order arrays saturate the objective function more rapidly. Geometry drift also decreases markedly with increasing *N* (*b* ≈ − 1.31, *R*^2^ ≈ = 0.77), reflecting increased geometric redundancy and reduced sensitivity to individual microphone perturbations. These trends suggest that additional microphones primarily stabilise the optimisation landscape rather than enabling proportionally better localisation performance.

Mean optimisation step size shows no reliable dependence on microphone count (CI spanning zero; *R*^2^ ≈ 0.03), indicating that the local move scale accepted by the optimiser is governed by geometric conditioning rather than sensor count alone.

Taken together, these results demonstrate a clear decoupling between optimisation behaviour and task-level performance. While higher microphone counts constrain the search space and promote rapid, stable convergence, they do not confer commensurate gains in volumetric localisation reliability. This finding reinforces the central conclusion of this study: carefully optimised low-order arrays can achieve high and robust localisation performance with substantially greater sensor economy than denser configurations.

### 3.5 Optimisation effects on array compactness

To assess whether improvements in localisation performance required physically larger arrays, I quantified the axis-aligned bounding-box dimensions of each configuration before and after optimisation (Table 2). For each run, array width, height, and depth were computed from microphone coordinates, and the maximum span across axes was used as a conservative measure of overall array extent.

**Table 1:** Power-law scaling of optimisation behaviour with microphone count. Each metric *y* was fitted with a model *y* = *a* N^*b*^ using log–log regression. Confidence intervals (CI) for *a* and *b* were estimated via bootstrap resampling (*B* = 2000). *R*^2^ is reported in log space.

| Metric | $n$ | $a$ | $b$ | $a_{CI,lo}$ | $a_{CI,hi}$ | $b_{CI,lo}$ | $b_{CI,hi}$ | $R^2_{log}$ |
| --- | --- | --- | --- | --- | --- | --- | --- | --- |
| Max. pass rate | 25 | 77.5 | 0.095 | 70.3 | 88.3 | 0.038 | 0.139 | 0.514 |
| Final pass rate | 25 | 77.5 | 0.095 | 70.3 | 88.4 | 0.038 | 0.138 | 0.514 |
| Convergence iter. | 19 | 220 | -2.10 | 51.3 | 1085 | -2.87 | -1.36 | 0.683 |
| Mean step size | 25 | 0.0105 | 0.195 | 0.0052 | 0.0222 | -0.16 | 0.53 | 0.032 |
| Drift RMS | 25 | 0.681 | -1.31 | 0.358 | 1.31 | -1.63 | -1.02 | 0.770 |

**Table 2:** Array compactness before and after optimisation. For each configuration (five runs each), median axis-aligned bounding-box spans (width *W*, height *H*, depth *D*; metres) are reported from microphone coordinates before (·_0_) and after (·_1_) optimisation. The maximum span `max` is the largest of (*W, H, D*) for a given state and provides a conservative measure of overall array extent. The ratio max_1_/max_0_ indicates whether optimisation expanded (> `1`) or compacted (< `1`) the array.

| ID | Configuration | $n$ | $W_0$ | $H_0$ | $D_0$ | $W_1$ | $H_1$ | $D_1$ | $\max_0$ | $\max_1$ | $\frac{\max_1}{\max_0}$ |
| --- | --- | --- | --- | --- | --- | --- | --- | --- | --- | --- | --- |
| 1 | Pyramid ($N_{\text{mic}}=4$) | 5 | 0.500 | 0.500 | 0.400 | 0.478 | 0.625 | 0.557 | 0.500 | 0.625 | 1.251 |
| 2 | Tetrahedron (4) | 5 | 0.491 | 0.491 | 0.491 | 0.620 | 0.707 | 0.529 | 0.491 | 0.707 | 1.441 |
| 3 | Octahedron (6) | 5 | 0.500 | 0.500 | 0.500 | 0.922 | 0.340 | 0.498 | 0.500 | 0.922 | 1.845 |
| 4 | Double tetrahedron (8) | 5 | 0.612 | 0.707 | 0.750 | 0.693 | 0.792 | 0.838 | 0.750 | 0.838 | 1.118 |
| 5 | Pentagonal antiprism (10) | 5 | 0.637 | 0.607 | 0.318 | 0.599 | 0.708 | 0.371 | 0.637 | 0.708 | 1.111 |
| 6 | Icosahedron (12) | 5 | 0.638 | 0.638 | 0.638 | 0.638 | 0.638 | 0.763 | 0.638 | 0.763 | 1.196 |
| 7 | Random (4) | 5 | 0.517 | 0.329 | 0.380 | 0.663 | 0.485 | 0.398 | 0.517 | 0.663 | 1.283 |
| 8 | Random (6) | 5 | 0.388 | 0.474 | 0.541 | 0.457 | 0.657 | 0.568 | 0.541 | 0.657 | 1.215 |
| 9 | Random (8) | 5 | 0.535 | 0.488 | 0.554 | 0.551 | 0.632 | 0.642 | 0.554 | 0.642 | 1.159 |
| 10 | Random (10) | 5 | 0.568 | 0.598 | 0.619 | 0.568 | 0.654 | 0.696 | 0.619 | 0.696 | 1.125 |
| 11 | Random (12) | 5 | 0.646 | 0.668 | 0.523 | 0.646 | 0.675 | 0.528 | 0.668 | 0.675 | 1.011 |

**Table 3:** Economy–responsiveness comparison of array geometries. Performance gain is defined as the absolute increase in volumetric pass rate following optimisation. Economy is quantified by the number of microphones *N*, and responsiveness is measured as gain per microphone (Δ/ `pass` *N*). Arrays are ranked primarily by positional economy (rank_pos_), with the angular economy shown for comparison. Higher responsiveness indicates a larger improvement in volumetric localisation reliability per sensor, highlighting configurations that benefit most from optimisation under fixed microphone budgets.

| ID | Geometry | $N$ | $\Delta_{\text{pos}}$ | $\Delta_{\text{pos}}/N$ | $\text{rank}_{\text{pos}}$ | $\Delta_{\text{ang}}$ | $\Delta_{\text{ang}}/N$ | $\text{rank}_{\text{ang}}$ |
| --- | --- | --- | --- | --- | --- | --- | --- | --- |
| 7 | Random | 4 | 52.60 | 13.15 | 1 | 26.17 | 6.54 | 1 |
| 1 | Pyramid | 4 | 42.60 | 10.65 | 2 | 9.69 | 2.42 | 2 |
| 2 | Tetrahedron | 4 | 29.46 | 7.36 | 3 | 8.97 | 2.24 | 3 |
| 3 | Octahedron | 6 | 43.11 | 7.19 | 4 | 0.00 | 0.00 | 8 |
| 8 | Random | 6 | 32.96 | 5.49 | 5 | 3.83 | 0.64 | 4 |
| 4 | Double tetrahedron | 8 | 10.51 | 1.31 | 6 | 0.00 | 0.00 | 6 |
| 9 | Random | 8 | 9.59 | 1.20 | 7 | 1.02 | 0.13 | 5 |
| 5 | Pentagonal antiprism | 10 | 11.48 | 1.15 | 8 | 0.00 | 0.00 | 9 |
| 10 | Random | 10 | 7.50 | 0.75 | 9 | 0.00 | 0.00 | 10 |
| 6 | Icosahedron | 12 | 8.67 | 0.72 | 10 | 0.00 | 0.00 | 7 |
| 11 | Random | 12 | 4.23 | 0.35 | 11 | 0.00 | 0.00 | 11 |

Across canonical geometries, optimisation generally resulted in moderate expansions of array size, though the magnitude and direction of change depended strongly on topology. Median max-span ratios ranged from 1.11 (pentagonal antiprism, 10 microphones) to 1.85 (octahedron, 6 microphones), corresponding to absolute increases of approximately 7–42 cm. The pyramidal and tetrahedral four-microphone configurations exhibited substantial expansion (ratios of 1.25 and 1.44, respectively), reflecting the need to escape geometrically constrained initial layouts. In contrast, higher-order canonical arrays showed comparatively restrained growth: the icosahedral configuration (12 microphones) increased only modestly (ratio = 1.20), despite its higher sensor count.

Random initialisations showed larger, more variable changes in compactness, particularly at low microphone counts. The random four-microphone configuration showed a median max-span ratio of 1.28, indicating pronounced expansion from initially compact or near-degenerate geometries. However, as the microphone count increased, random arrays exhibited progressively smaller relative changes, with the random 12-microphone configuration remaining almost unchanged after optimisation (ratio = 1.01). This pattern indicates that higher-order arrays inherently provide sufficient geometric flexibility to accommodate optimisation without substantial physical expansion.

Importantly, no monotonic relationship was observed between microphone count and final array size. Optimised arrays with 8–12 microphones occupied comparable spatial extents despite differing sensor counts, reinforcing the conclusion that improved performance at higher *N* arises primarily from geometric redundancy rather than increased physical footprint. These results demonstrate that substantial gains in volumetric localisation reliability can be achieved without proportional increases in array size, supporting the feasibility of compact, portable array designs for field deployment.

## 4 DISCUSSION

This study introduces a task-oriented framework for microphone-array optimisation that explicitly links array geometry to volumetric localisation reliability under realistic field constraints. The central finding challenges a common assumption in acoustic array design: increasing microphone count does not guarantee proportional improvements in localisation performance. Instead, carefully optimised low-order arrays—particularly four-to-six-microphone configurations—can achieve comparable or superior performance to higher-order systems when evaluated holistically through metrics of economy and responsiveness to optimisation.

This result has direct practical relevance for bioacoustic field studies. In deployments where portability, setup speed, and mechanical robustness are paramount, optimised four-microphone arrays offer a favourable balance between accuracy and deployability. When normalised by sensor count, four-microphone configurations consistently exhibit the highest performance gain per microphone, making them particularly efficient for near to mid-field volumetric localisation. These findings motivate a re-evaluation of common array-design heuristics in bat bioacoustics, where larger arrays are often adopted by default without systematic assessment of whether their additional complexity yields proportional gains in usable sensing volume.

In many acoustic-tracking studies, localisation is appropriately used as an intermediate measurement for reconstructing behaviour rather than evaluated as a methodological outcome in its own right (de Framond et al., 2023; Hiryu et al., 2010; Holderied & von Helversen, 2003; Jakobsen et al., 2025; Lewanzik & Goerlitz, 2018). Consequently, spatial uncertainty may not always be reported in detail or evaluated relative to the behavioural quantities subsequently derived from the reconstructed positions. This highlights the importance of matching localisation accuracy to the spatial and temporal scale of the question being addressed.

The implications of localisation uncertainties are non-trivial. For example, a positional error of 0.25–0.5 m over a 50 ms interval—comparable to a typical inter-pulse interval during search or approach—corresponds to a velocity uncertainty of approximately 5–10 m s^−1^. This range overlaps the flight velocities reported for aerial-hawking vespertilionid bats, including *Pipistrellus* and *Myotis* species (Haron et al., 2026; Jakobsen et al., 2025), implying that biologically meaningful kinematic signatures can be obscured by localisation error alone. Although extreme errors may be identifiable—for example, an apparent velocity change from 2 to 20 m s^−1^ over 50 ms implies a physiologically implausible acceleration exceeding 36 *g*—less conspicuous errors may still affect behavioural inference. Localisation uncertainty should therefore be evaluated relative to the spatial and temporal scale of the behaviour under investigation.

Furthermore, visualisation or smoothing of reconstructed flight paths can make the influence of positional uncertainty less apparent. Although many studies address this by restricting the analysis volume or excluding outliers, task-specific evaluation of array geometry provides an additional means of determining whether a configuration can resolve the behavioural features under investigation. The present framework addresses this need by enabling microphone arrays to be designed and evaluated against explicit accuracy requirements.

This study extends the Widefield Acoustics Heuristic (Umadi, 2025c) by inverting its original forward perspective. Whereas the earlier framework characterised the accuracy produced by a given array geometry, the present work uses that characterisation as an objective function to iteratively refine the geometry to achieve a desired field of accuracy. In this sense, the approach transforms array design from a descriptive exercise into a constructive one.

The optimisation and evaluation were restricted to forward-facing upper octants, reflecting typical field deployments where arrays are oriented toward expected flight paths rather than enclosing the source volume isotropically. This restriction improves ecological relevance while reducing computational cost. Given a grid resolution of 0.25 m, 55 optimisation trajectories, 25 iterations per run, and 11 configurations, expanding the evaluated volume would substantially increase computational expense without altering the underlying optimisation logic. Importantly, the algorithm itself is agnostic to the chosen evaluation field: with sufficient computational budget, the same framework can be applied to alternative spatial constraints, target volumes, or task-specific accuracy thresholds.

The optimisation problem addressed here is inherently geometric. Localisation error in TDOA-based systems arises from the interaction between timing uncertainty and sensor geometry, commonly expressed through geometric dilution of precision (GDOP) (Bancroft, 1985; Chan & Ho, 1994). Even under ideal timing conditions, array geometry imposes structural limits on the regions of space that can be resolved accurately. Crucially, these limits are highly non-uniform and depend on both array topology and source position.

The results demonstrate that while higher microphone counts improve raw spatial sampling density, the gains saturate rapidly once geometric constraints dominate error structure (Figs. 4 & 5). The observed convergence behaviour (Fig. 2) shows persistent configuration-dependent differences under the specified constraints and computational budget. Adding microphones within a fixed aperture does not fundamentally overcome distance-dependent degradation in localisation accuracy (Fig. 5), reinforcing the notion that geometry, rather than sensor count alone, governs usable sensing volume under the tested conditions.

### 4.1 Geometric comparison of gained performance

To disentangle geometric constraints from optimisation artefacts, I evaluated six canonical three-dimensional geometries—a pyramid, tetrahedron, octahedron, double tetrahedron, pentagonal antiprism, and icosahedron—alongside randomly initialised arrays with matched microphone counts. Canonical geometries are attractive in practice because they are mechanically simple to construct, often benefit from symmetry, and can be realised using standard structural components such as commercially available joints and connectors. However, such geometries could, in principle, act as restrictive initialisation points that bias optimisation toward local optima.

Random initialisations address this concern by sampling the configuration space stochastically, providing a broader representation of possible geometries. Because the space of valid microphone configurations is effectively continuous and infinite, both canonical and random configurations should be interpreted as sparse samples from an underlying high-dimensional design space. Repeated optimisation from both structured and random initial conditions helps distinguish persistent configuration-dependent performance patterns from optimisation path-dependence (Fig. 4). Convergence of both initialisation strategies toward similar performance levels indicates that the observed patterns are robust under the present constraints and search budget, although it does not establish a global geometric optimum.

The optimised arrays need not converge on a single ideal geometry. From both canonical and random starting configurations, the optimiser made goal-oriented adjustments to microphone positions that reduced poorly constrained regions within the target volume by improving the spatial diversity of the time-difference measurements. This process is analogous to adjusting fingertip positions to obtain a stable grasp: several distinct arrangements can provide an effective solution. The resulting arrays should therefore be interpreted as alternative locally optimised configurations for the specified observation volume, accuracy criterion, and physical constraints, rather than as approximations to a universally optimal shape.

### 4.2 Microphone economy and performance scaling

The economy–responsiveness analysis provides a quantitative basis for ranking array geometries in terms of performance gain per microphone (Table 3). Across both canonical and random configurations, four-microphone arrays consistently rank highest in positional economy, achieving the largest absolute increases in volumetric pass rate when normalised by sensor count. While higher-order arrays often achieve slightly higher absolute pass rates, the marginal benefit per additional microphone decreases sharply beyond *N* = 4 − 6.

This pattern is reinforced by the power-law scaling analysis (Table 1). Maximum and final volumetric pass rate show weak positive scaling with microphone count: their bootstrap confidence intervals exclude zero, but the shallow exponent (*b* ≈ 0.095) indicates only a slow increase. In contrast, optimisation dynamics show strong and systematic scaling with *N*: the convergence iteration count and overall geometric drift both decrease approximately as *N*^−2^ and *N*^−1^, respectively. These results indicate that increasing array order primarily constrains the optimiser’s degrees of freedom, rather than yielding proportional improvements in usable localisation volume.

At the same time, denser arrays confer increased robustness to structural perturbations. With more microphones, small deviations in individual sensor positions exert a reduced influence on the global solution, effectively smoothing the error landscape. Under conditions where additional geometric freedom is available—such as larger allowable aperture sizes, extended arm lengths, or actively adjustable mounting geometries—higher-order arrays may therefore access substantially improved accuracy profiles (Benesty et al., 2008; Brandstein & Ward, 2001). In such cases, the benefits of increased redundancy and spatial sampling may outweigh the diminishing returns, highlighting that the optimal microphone count is ultimately application-dependent.

In field settings where the primary objective is reliable near to mid-field localisation—such as reconstructing flight trajectories, estimating spatial decision rules, or resolving interactions among nearby individuals—optimised four-microphone arrays therefore offer a favourable balance between accuracy, responsiveness to optimisation, and economy. The modest and saturating gains achieved by denser arrays suggest that additional sensors are often better traded for improvements in timing precision, deployment geometry, or experimental flexibility, rather than increased microphone count alone.

### 4.3 Accuracy: task-specific design considerations

Positional accuracy was treated as the primary optimisation objective, reflecting its central role in reconstructing three-dimensional trajectories in the near field. Angular accuracy was evaluated as a secondary metric, providing complementary information about directional resolution. As expected, angular accuracy improved with increasing microphone count (Table 3), reflecting the increased directional diversity afforded by denser arrays.

However, the economic analysis reveals that angular gains per microphone remain lower for higher-order configurations. Consequently, the choice of microphone count should be guided by task requirements: applications prioritising precise directional cues (e.g., beamforming) may justify denser arrays, whereas studies focused on spatial reconstruction within a limited volume benefit more from optimised low-order geometries (Van Trees, 2002). The angular error distributions shown in Figures 5 and 6 further demonstrate that denser arrays yield more robust and consistently low angular errors across distance, making them particularly well suited for far-field applications such as source separation and directional filtering (Benesty et al., 2008; Brandstein & Ward, 2001), where angular stability rather than absolute positional accuracy is the primary constraint.

Beyond localisation accuracy, optimised microphone-array geometry has direct implications for acoustic-source separation in conservation bioacoustics, where emerging techniques are actively addressing the problem at individual to population, and niche levels (Blumstein et al., 2011; Izadi et al., 2020; Lin & Tsao, 2020; Rhinehart et al., 2020; Tolkova & Klinck, 2022). In dense acoustic environments, overlapping calls from multiple individuals or species constitute a major limitation for abundance estimation and behavioural inference. Arrays that yield poorly resolved spatial cues yield ambiguous direction estimates and unstable separation, even at high signal-to-noise ratios. By maximising volumetric localisation reliability under realistic constraints, the proposed optimisation framework implicitly improves the spatial separability of acoustic sources throughout the observation volume. This establishes a natural extension of the method toward array designs tailored not only for localisation, but also for multi-source disentanglement in conservation monitoring contexts, thus directly contributing to ongoing developments (Blumstein et al., 2011; Laiolo, 2010; Sugai et al., 2019).

### 4.4 Build and deployment considerations

An important practical insight is that relatively small geometric perturbations can produce measurable changes in localisation performance (Fig. 7). This sensitivity implies that unintentional deviations during construction or deployment—such as slight arm misalignments or mounting asymmetries—can materially affect array performance. Consequently, careful attention to mechanical design and reproducibility is essential, particularly for arrays operating near the threshold of acceptable accuracy.

At the same time, the finding that optimised four-microphone arrays achieve very high performance opens new possibilities for reusable, adaptable array systems. Compact hardware platforms such as Batsy4-Pro (Umadi, 2025b) can be paired with adjustable or telescopic arm structures mounted on ball joints, allowing rapid reconfiguration for task-specific optimisation. This combination enables a single system to be repurposed across laboratory and field contexts, enhancing both logistical efficiency and experimental control.

### 4.5 Broader applicability and limitations

The WAH-*i* algorithm can be applied to any acoustic signal for which reliable time differences of arrival can be estimated. The underlying conclusion that localisation reliability depends strongly on array geometry should therefore extend to other bioacoustic and engineered systems. The numerical results reported here, however, are conditional on the values of the simulation parameters and pertain to the specified observation volume and physical constraints. Signal bandwidth, centre frequency, duration, directionality, and signal-to-noise ratio affect delay-estimation precision and may consequently alter absolute errors, attainable pass rates, configuration rankings, and the geometry favoured for a particular task. In WAH-*i*, signal type, upper and lower frequencies, duration, sampling rate, and signal-to-noise ratio are user-configurable, allowing these conditions to be matched to a particular species or recording context. Application to birds, amphibians, other bat call types, or different recording media and environments therefore requires rerunning the optimisation with representative signal and noise parameters rather than transferring the present numerical rankings directly.

The present simulations further assume a stationary source, homogeneous sound speed, no multipath reflections, and perfect sensor synchrony (Umadi, 2025a, 2025c). Although the software can generate constant-frequency (CF) signals, comparable performance cannot be assumed because narrowband signals may yield less precise or ambiguous correlation peaks. Source motion introduces an additional distinction: CF signals are particularly susceptible to systematic localisation errors under relative motion, whereas short FM calls can retain robust median accuracy while showing broader upper-tail errors (Umadi, 2025b, 2025c). These controlled assumptions isolate geometry-dependent effects, but field-specific propagation, noise, motion, and synchronisation should be incorporated when designing an array for deployment.

Computational demand also limits the resolution and breadth of the present search. Each candidate microphone displacement requires a full volumetric evaluation, so cost increases approximately with the inverse cube of grid spacing for a fixed three-dimensional volume. More stringent accuracy or pass-rate requirements can prolong convergence, increase stalls and stochastic shakes, and require larger iteration budgets and more independent runs. Because the optimiser searches locally around each initial configuration, it does not guarantee a global optimum; the observed plateaus therefore describe performance under the specified objective, stopping rule, and computational budget rather than absolute geometric limits.

Several extensions could reduce these constraints or broaden the framework’s use. Optimisation histories provide structured training data for surrogate models (Forrester et al., 2008), including regression or lightweight neural-network approaches (Lee et al., 2021), that could identify promising regions of configuration space before refinement by more computationally demanding searches (Raissi et al., 2019; Shahriari et al., 2016). The same task-level objective could also support adaptive or reconfigurable arrays whose geometry responds to changing source regions, environmental conditions, or auxiliary information from inertial, visual, radar, or depth sensors.

The general design principle—balancing spatial discrimination, robustness, coverage, and sensor count—may also be useful beyond bioacoustics. Multi-microphone hearing-assistive devices, for example, use array processing to enhance spatial selectivity, suppress interference, and track target speakers (Doclo et al., 2010; Thiemann et al., 2016; Widrow & Luo, 2003). Although these systems operate at different spatial scales, optimisation-driven geometry selection could complement emerging learning-based approaches to spatial hearing and direction-of-arrival estimation (He et al., 2018; Zhang et al., 2021). These applications require specific signal and propagation models, but they illustrate how the algorithm can be transferred without assuming that the bat-specific configurations or performance values reported here remain optimal.

Taken together, these findings show that microphone-array design is best treated not as a choice among conventional geometries or simply as a question of sensor number, but as a goal-specific optimisation problem defined by explicit spatial, acoustic, and physical requirements. WAH-*i* provides a practical means of translating those requirements into deployable array configurations, enabling localisation performance to be evaluated—and geometry refined—before behavioural inference or field deployment.

## ACKNOWLEDGEMENTS

I thank Uwe Firzlaff and colleagues at the TUM for their support and institutional assistance throughout the development of this work. I also extend my sincere thanks to the three reviewers for their careful engagement with the manuscript and their constructive feedback, and to the editors for their thoughtful handling and careful evaluation.

## AUTHOR CONTRIBUTIONS

This study was conceived and executed by the sole author.

## ETHICAL STATEMENT

This study did not involve animal handling, experimentation, or data collection from live animals.

## DATA & CODE AVAILABILITY

All data and code used in this study, as well as the WAH-*i* installable packages, are archived on Zenodo at https://doi.org/10.5281/zenodo.21534092.

User-guide and the documentation for the WAH-*i* software are published at https://raviumadi.github.io/WAHi-Optimiser.

## SUPPLEMENTARY INFORMATION

### A PERFORMANCE GAIN TABLES

#### A.1 Positional Performance

**Table A1:** Positional error statistics before and after optimisation. Pass rate denotes the percentage of grid points with positional error ≤ 15 cm. Error statistics (mean, median, interquartile range (IQR), and 95th percentile (p95)) are reported in centimetres.

| Configuration | $N$ | Pass<sub>0</sub><br>(%) | Mean<sub>0</sub><br>(cm) | Median<sub>0</sub><br>(cm) | IQR<sub>0</sub><br>(cm) | p95<sub>0</sub><br>(cm) | Pass<sub>1</sub><br>(%) | Mean<sub>1</sub><br>(cm) | Median<sub>1</sub><br>(cm) | IQR<sub>1</sub><br>(cm) | p95<sub>1</sub><br>(cm) |
| --- | --- | --- | --- | --- | --- | --- | --- | --- | --- | --- | --- |
| Pyramid | 4 | 54.1 | 32.7 | 11.6 | 34.6 | 136.8 | 96.7 | 5.1 | 3.8 | 6.1 | 13.6 |
| Tetrahedron | 4 | 52.7 | 31.0 | 13.0 | 38.0 | 118.3 | 82.1 | 15.9 | 4.2 | 9.6 | 104.1 |
| Octahedron | 6 | 52.0 | 93.1 | 13.8 | 30.5 | 84.9 | 95.2 | 6.0 | 4.8 | 6.5 | 15.0 |
| Double tetrahedron | 8 | 84.7 | 8.3 | 6.0 | 8.8 | 24.2 | 95.2 | 5.3 | 4.0 | 5.9 | 14.9 |
| Pentagonal antiprism | 10 | 83.7 | 7.6 | 5.2 | 9.1 | 21.5 | 95.2 | 5.3 | 3.8 | 5.5 | 14.8 |
| Icosahedron | 12 | 87.5 | 7.1 | 5.2 | 7.7 | 18.6 | 96.2 | 4.9 | 3.5 | 4.7 | 14.4 |
| Random | 4 | 33.9 | 2608.9 | 36.3 | 104.6 | 587.5 | 86.5 | 11.1 | 4.9 | 8.5 | 26.4 |
| Random | 6 | 62.3 | 518.4 | 10.4 | 20.6 | 99.0 | 95.3 | 5.7 | 4.0 | 5.8 | 14.9 |
| Random | 8 | 85.7 | 9.2 | 5.3 | 8.4 | 25.5 | 95.3 | 5.5 | 4.0 | 5.8 | 14.6 |
| Random | 10 | 88.5 | 7.1 | 5.0 | 7.6 | 21.2 | 96.0 | 5.2 | 3.8 | 5.5 | 14.2 |
| Random | 12 | 92.7 | 5.9 | 4.1 | 6.3 | 17.3 | 96.9 | 5.0 | 3.6 | 5.1 | 13.4 |

#### A.2 Angular Performance

**Table A2:** Angular localisation error statistics before and after optimisation. Pass rate denotes the percentage of grid points with angular error ≤ 15^°^. Angular error statistics (mean, median, interquartile range (IQR), and 95th percentile (p95)) are reported in degrees.

| Configuration | $N$ | Pass<sub>0</sub><br>(%) | Mean<sub>0</sub><br>(deg) | Median<sub>0</sub><br>(deg) | IQR<sub>0</sub><br>(deg) | p95<sub>0</sub><br>(deg) | Pass<sub>1</sub><br>(%) | Mean<sub>1</sub><br>(deg) | Median<sub>1</sub><br>(deg) | IQR<sub>1</sub><br>(deg) | p95<sub>1</sub><br>(deg) |
| --- | --- | --- | --- | --- | --- | --- | --- | --- | --- | --- | --- |
| Pyramid | 4 | 90.3 | 4.02 | 0.97 | 2.14 | 24.6 | 100.0 | 0.72 | 0.45 | 0.59 | 2.26 |
| Tetrahedron | 4 | 82.1 | 13.4 | 1.46 | 5.24 | 75.7 | 91.1 | 6.90 | 0.64 | 1.39 | 52.5 |
| Octahedron | 6 | 100.0 | 1.91 | 1.22 | 1.79 | 6.78 | 100.0 | 0.98 | 0.88 | 0.77 | 2.32 |
| Double tetrahedron | 8 | 100.0 | 0.76 | 0.69 | 0.62 | 1.76 | 100.0 | 0.63 | 0.55 | 0.51 | 1.37 |
| Pentagonal antiprism | 10 | 100.0 | 1.04 | 0.72 | 0.79 | 3.24 | 100.0 | 0.59 | 0.47 | 0.45 | 1.48 |
| Icosahedron | 12 | 100.0 | 0.66 | 0.57 | 0.54 | 1.43 | 100.0 | 0.55 | 0.46 | 0.38 | 1.24 |
| Random | 4 | 73.8 | 20.2 | 3.63 | 15.2 | 136.2 | 99.9 | 0.98 | 0.68 | 1.14 | 2.69 |
| Random | 6 | 96.2 | 2.90 | 0.97 | 1.25 | 7.67 | 100.0 | 0.73 | 0.58 | 0.62 | 1.85 |
| Random | 8 | 98.9 | 1.05 | 0.58 | 0.62 | 1.85 | 99.9 | 0.60 | 0.49 | 0.47 | 1.35 |
| Random | 10 | 100.0 | 0.64 | 0.52 | 0.52 | 1.51 | 100.0 | 0.56 | 0.47 | 0.43 | 1.35 |
| Random | 12 | 100.0 | 0.54 | 0.44 | 0.45 | 1.31 | 100.0 | 0.48 | 0.39 | 0.37 | 1.15 |

### B WAH-*I* RUNS IMPLEMENTATION

The following sections describe the implementation and analysis procedures with explicit reference to variable names and structures used in the accompanying codebase; all such variables are defined and documented within the released source code.

#### B.1 Batch experimental design and run generation

The batch runner (`run_iwah_experiment_batch.m`) executed optimisation runs under two start-condition classes:

1. **CSV-defined canonical geometries (“CSV starts”)**. Six standard geometries were provided as CSV files (`configs/*. csv`), each defining an initial array layout in Cartesian coordinates:
  - 4-microphone Pyramid (`4mics_Pyramid.csv`)
  - 4-microphone Tetrahedron (`4mics_Tetrahedron_small.csv`)
  - 6-microphone Octahedron (`6mics_Octahedron.csv`)
  - 8-microphone Double tetrahedron (`8mics_double_tetrahedorn_small.csv`)
  - 10-microphone Pentagonal antiprism (`10mics_pentagonal_antiprism.csv`)
  - 12-microphone Icosahedron (`12mics_icosahedron.csv`)
  If enabled (`EXP.useCsvNativeN=true`), the microphone count *N*_mic_ was read directly from the number of rows in the CSV file; otherwise, *N*_mic_ could be overwritten by the batch settings.
2. **Random-start geometries (“random starts”)**. For each *N*_mic_ ∈ {4, 6, 8, 10, 12}, a random initial geometry was generated internally by the optimiser (see below) and then refined.

For each condition (canonical or random) I ran *k* = 5 independent repeats (`EXP.repeats=5`). Unless debugging, runs were executed in parallel using MATLAB’s `parfor` (`EXP.useParfor=true`) with a local pool (`parpool`).

##### B.1.1 Simulation space and test grid

All optimisation runs were evaluated on a common three-dimensional localisation test grid defined by:

- outer radius *R* = 2.0 m (`P.R`)
- inner exclusion radius *R*_in_ = 0.30 m (`P.Rin`)
- grid step *s* = 0.250 m (`P.step`)

The grid thus sampled candidate source locations in a volumetric shell around the array centre (origin). For each iteration, the optimiser computed localisation errors at these grid points (see below) and stored the grid points as `iter(k).` `gridPts`.

##### B.1.2 Geometric constraints and feasibility

To prevent degenerate geometries and to enforce practical build constraints, candidate microphone layouts were constrained as follows:

- **Bounded array extent:** all microphone positions were constrained within a sphere of radius ***L***_max_ = 0.50 m (`P. Lmax`).
- **Minimum inter-microphone distance:** microphone pairwise distances were constrained to exceed `P.minPairDist` (=6.25 cm).
- **Minimum source-to-microphone distance:** grid points too close to any microphone were excluded using `P.minDistToAnyMic`=0.08 m, with a soft exclusion parameter `P.epsDist`=0.03 m.

For CSV-defined geometries, the batch pipeline recentered the array coordinates to the origin (`CSV.recenter=true`) and optionally repaired minor constraint violations (`CSV.autoRepair=true`). The repair procedure attempted up to 200 iterations (`CSV.repairMaxTries`) with a jitter perturbation that increased multiplicatively (`CSV.repairJitter 0=1e-3`,growth `CSV.repairJitterGrowth=1.15`). If enabled, repair clamped extents to ***L***_max_ (`CSV.repairClampLmax=true`) andscaling adjustments (`CSV.repairPreferScale=true`).

##### B.1.3 Acoustic signal and propagation parameters

Each localisation evaluation was based on a simulated biosonar-like call, parameterised by:

- sampling rate *f*_*s*_ = 192 kHz (`P.fs`)
- call duration *d* = 5 ms (`P.d`)
- FM sweep from *f*_0_ = 30 kHz to *f*_1_ = 70 kHz (`P.f0`, `P.f1`)
- additive noise level `P.snr_db`=40 dB
- tail window parameter `P.tail`=200 (used internally by the call generator/processing)
- call type `P.callType=`‘`FM`’
- source velocity v = [0, 0, 0] (`P.velocity`), i.e. stationary source
- speed of sound *c* = 343 m s^−1^ (P`.c`)

These parameters were held constant across conditions and repeats to isolate the effect of geometry and microphone count.

##### B.1.4 Optimisation objective, pass criterion, and penalties

At each iteration, the optimiser produced localisation error statistics over the test grid and computed:

- **Position error** *e*_pos_ in cm (`stats.err_cm`), and/or
- **Angular error** *e*_ang_ in degrees (`stats.ang_deg`)

depending on the selected evaluation metric.

A grid point was deemed a *pass* if its position error satisfied

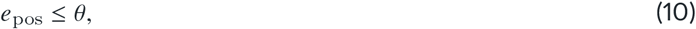

where the pass threshold was defined as `P.threshold_cm` with

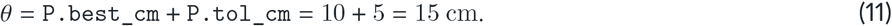

The *pass rate* (`stats.passRate`) was the fraction of grid points passing the criterion.

To discourage pathological failures, runs used a large penalty for failing or invalid points (`P.failPenalty_cm`=1000). Additionally, the optimiser targeted a desired pass rate `P.targetPass`=0.95 and used an internal weighting parameter `P.lambdaMean`=0.15 to balance mean error and pass-based objectives (exact internal objective combination is implemented within `iwah_run_optimiser_core` and recorded as `iter(k).J`).

##### B.1.5 Optimiser configuration and stopping behaviour

Optimisation used the following control parameters (`OPT`):

- maximum iterations: 25 (`OPT.maxIters`)
- probing density: 14 probes per microphone per iteration (`OPT.probesPerMic`)
- initial step fraction: 0.25 (`OPT.initStepFrac`)
- minimum step fraction: 0.03 (`OPT.minStepFrac`)
- stall limit: 12 (`OPT.stallLimit`)
- “shake” fraction: 0.08 (`OPT.shakeFrac`)
- improvement tolerance: `OPT.improveEpsJ`=10^−6^

The optimiser recorded intermediate shake steps using non-integer iteration indices (e.g. *it* = 0.5), which were later excluded from trajectory summaries to align runs on integer iteration counts.

##### B.1.6 Octant selection

The experiment allowed restricting evaluations to specific spatial octants (`Q`). In the current batch settings, only two regions were enabled: `frontTopLeft=true` and `frontTopRight=true`, while all others were disabled. This restriction was applied consistently across all conditions and repeats.

##### B.1.7 Random seeds and reproducibility

Each run used a unique per-run seed generated without fixed seeding to avoid correlations across parallel workers. The seed combined a high-resolution time component with the process ID and an additional random draw, then reduced modulo 2^31^ − 1. Each run stored its seed (argument `seed`) within the run history and metadata for reproducibility.

##### B.1.8 Run outputs and file naming

Each optimisation produced two outputs:

- `H`: the full history structure containing iteration-wise geometry (`iter(k).M`), objective (`iter(k).J`), statistics (`iter(k).stats`), and grid points (`iter(k).gridPts`), as well as start/final metadata in `H.meta`.
- `runInfo`: a lightweight struct with identifiers (configuration tag, repeat index, start mode, CSV stem if applicable).

Files were saved as individual `.mat` files to `runs/` with filenames encoding condition, final microphone count, repeat index, and timestamp.

#### B.2 Analysis pipeline and figure generation

All batch runs were analysed using `iwah_analyse_batch_runs.m`, which loads all `.mat` files in `runs/` and extracts common summary products. Unless stated otherwise, I report **position error** (`METRIC=`‘`pos`’) in cm. Below I describe each analysis component in the same order as the script, and I indicate the corresponding figures produced by the pipeline.

##### B.2.1 Run ingestion, grouping, and “before” vs “after” selection

For each run file, I identified:

- microphone count *N*_mic_ as the number of rows in `H.meta.final.M` (fallback: `H.meta.start.M`);
- start type (`csv` vs `random`) using `H.meta.start.csvPath` (or fallback string parsing of `H.meta.start.source`);
- condition key as the CSV filename stem for CSV starts and as `randomStart` for random starts; and
- a short label for each condition using a fixed mapping (`COND_LABELS`).

I defined **“before”** as the first recorded iteration (`iBase=1`). I defined **“after”** using one of two modes:

1. `AFTER_MODE=`‘`final`’: the last recorded iteration (`iFinal=numel(H.iter)`), or
2. `AFTER_MODE=`‘`best`’: the iteration achieving the maximum pass rate, with ties broken by (i) smaller *p*_95_ error and then (ii) larger objective value *J*.

###### Masking of unused test points

Errors were optionally computed only over valid/used test points (`IGNORE_UNUSED=true`) by applying the boolean vector `stats.usedMask`. This step excludes grid points that the localisation/evaluation routine marks as unused (e.g. invalid due to constraints or internal screening) and prevents unused points from biasing histograms and summary statistics.

##### B.2.2 Figure: pass-rate trajectories across iterations

This figure summarises optimisation dynamics using pass-rate trajectories. For each *N*_mic_ ∈ {4, 6, 8, 10, 12} I grouped runs by condition (CSV geometry or random start) and computed a robust central trajectory:

1. I extracted iteration-wise pass rates `iter(k).stats.passRate`.
2. Only **integer** iteration indices were retained to exclude shake steps (e.g. *it* = 0.5) and to ensure consistent alignment.
3. Each run trajectory was aligned onto a common integer iteration axis by forward-filling missing iterations (holding the last observed value).
4. I summarised across repeats using the median trajectory and a shaded uncertainty band, using either the interquartile range (IQR: 25–75%) or a 10–90% band (parameter `TRAJ.band`).

##### B.2.3 Figure: “average-run” error distributions before vs after

This figure visualises how optimisation redistributes localisation error. For each *N*_mic_ and start type (CSV vs random), I computed filled histograms for pre- and post-optimisation errors:

1. For each run, I extracted the error vector `stats.err_cm` (or `stats.ang_deg`) at the “before” and “after” iterations.
2. If enabled, I filtered these vectors using `stats.usedMask`.
3. To avoid histogram domination by extreme failures, I optionally capped errors at a multiple of the pass threshold (`HIST.capMode`: `2xThreshold` or `4xThreshold`).
4. Within each panel, histogram bin edges were determined from the pooled (capped) values to ensure a consistent binning scheme across “before” and “after”.
5. The displayed histogram is an *average-run* distribution: for each run I computed a probability histogram (normalised to sum to 1), then averaged these probabilities across repeats.

A special case was applied for *N*_mic_ = 4 CSV starts: Pyramid and Tetrahedron conditions were plotted separately (distinct colours) to allow direct comparison of the two 4-microphone canonical geometries within the same panel.

##### B.2.4 Figure: accuracy vs distance from array centre

This figure quantifies spatial structure in localisation error by summarising error as a function of radial distance from the array centre. For each run I computed the radial coordinate *r* = ∥x∥_2_ for each grid point `gridPts`. I then:

1. defined *n*_*r*_ = 10 radial bins spanning the range of *r* values in the dataset;
2. within each bin, computed robust statistics for “before” and “after” errors: median and a quantile band (default 10–90%, parameters `RAD.qLo`, `RAD.qHi`);
3. optionally applied an error cap (default `RAD.capForScatter=`‘`2xThreshold`’) to reduce the influence of catastrophic failures on the quantile summaries; and
4. aggregated across repeats by taking the median of run-wise medians (and of run-wise quantiles) at each radial bin centre.

##### B.2.5 Evaluation of before–after differences and robust statistics

Across all analyses, pre- and post-optimisation differences were evaluated using robust, distribution-aware summary statistics computed on masked error vectors. For each run, error values were first filtered using the boolean vector `stats.usedMask`, thereby excluding grid points that were deemed invalid or unused by the localisation routine (e.g. due to geometric infeasibility or internal screening). All reported statistics are therefore conditioned on the same set of valid test points before and after optimisation.

To characterise localisation performance beyond the mean, I focused on upper-tail behaviour using the 95th percentile of the error distribution (*p*_95_). This choice reflects worst-case but non-pathological performance and avoids sensitivity to extreme outliers, which are handled separately (see below). For each configuration group, I computed:

- pass rate (fraction of points with *e* ≤ *θ*),
- mean error,
- median error,
- interquartile range (IQR; 75th–25th percentile), and
- *p*_95_ error.

These quantities were computed for both “before” and “after” states, and differences were defined as Δ*x* = *x*_after_ − *x*_before_.

##### B.2.6 Figure: representative geometry and error-field visualisation

This figure provides a qualitative illustration of how optimisation alters both microphone geometry and the resulting spatial error field. For a selected canonical configuration (specified by `SELECT_GEOM`), I identified a representative run as the one achieving the highest post-optimisation pass rate, with ties broken by the smallest *p*_95_ error. For this run, I visualised:

- the microphone coordinates at the initial state and after optimisation,
- the three-dimensional test grid coloured by localisation error,
- the outer evaluation sphere (*R*) and maximum geometry extent (***L***_max_), and
- the array centre (origin).

Error values were capped at 2× the pass threshold to stabilise colour scaling and highlight spatial structure in the bulk of the distribution. Only grid points marked as valid by `usedMask` were included. This figure serves as an intuitive complement to the statistical summaries presented elsewhere.

##### B.2.7 Pooled summary statistics table

In addition to the figures, I computed pooled summary statistics across runs for each configuration group. For CSV starts, statistics were stratified by canonical geometry and microphone count; for random starts, statistics were stratified by microphone count only. Error vectors were pooled across all repeats within each group after applying `usedMask`.

For each group, I report pass rate, mean, median, IQR, and *p*_95_ error for both pre- and post-optimisation states. These pooled values form the basis for quantitative comparisons across configurations and array sizes.

##### B.2.8 Delta mosaic of optimisation effects

To compactly summarise the direction and magnitude of optimisation effects across all configurations, I constructed a *delta mosaic* heatmap. Each row corresponds to a configuration (canonical geometries and random starts), and each column corresponds to a summary metric (pass rate, mean, median, IQR, *p*_95_, and optional tail fraction).

For interpretability, deltas were displayed such that positive values indicate improvement. Specifically, pass-rate deltas were used directly (Δ in percentage points), whereas error-based deltas were sign-inverted so that reductions in error appear as positive values.

Colour scaling was made robust by setting symmetric limits based on the 95th percentile of |Δ| across all cells. This prevents a small number of extreme changes from dominating the visual encoding while preserving relative differences among configurations.

##### B.2.9 Tail behaviour and catastrophic errors

To explicitly track severe localisation failures, I quantified the fraction of grid points with errors exceeding a multiple of the pass threshold. Errors larger than 4 × *θ* were classified as catastrophic. For each configuration group, I computed the percentage of such tail points before and after optimisation and included the corresponding change (Δ tail fraction) as an optional column in the delta mosaic.

This separation allows improvements in the bulk of the distribution (median, IQR) to be interpreted independently from changes in extreme failure rates.

##### B.2.10 Optimisation-induced geometry drift

To quantify how much the optimiser altered microphone positions, I analysed geometry drift using `iwah_geometry_ drift_plots_etc.m`. For each run, I extracted start and end microphone coordinates and computed per-microphone displacement vectors

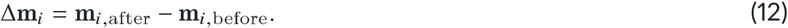

Drifts were pooled across microphones and repeats within each configuration and visualised as boxplots for the *x*, y, and *z* components separately. A horizontal reference line at zero indicates no net displacement.

This analysis allows assessment of whether performance gains arise from large geometric rearrangements or from comparatively subtle adjustments.

##### B.2.11 Runtime diagnostics for pass-rate discrepancies

To explain why different start geometries can exhibit different *achievable* maximum pass rates (and not merely different convergence speeds), I computed a set of per-run runtime diagnostics from the full optimiser history (`H.iter`) and summarised these by configuration. Diagnostics were extracted from integer iteration indices only (i.e. excluding non-integer “shake” steps) to match the trajectory alignment used in Figure 1.

For each run, I derived the pass-rate trajectory {*p*(i*t*)} from `iter(k).stats.passRate`. If the stored values were fractions (0–1), they were converted to percentages for reporting. I then computed:

- **Achievable max pass rate:** *p*_max_ = max_*it*_ *p* (*it*).
- **Final pass rate:** *p*_final_ = *p* (i*t*_end_), and **net change** Δ*p* = *p*_final_ − *p* (*it*=1).
- **Convergence iteration:** the first iteration at which the run reached a fixed fraction of its own maximum,

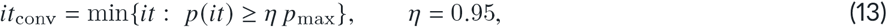

thereby separating differences in *attainable* performance from differences in *speed* of improvement.
- **Step-size proxy:** a per-iteration geometry update magnitude computed from successive microphone-coordinate matrices,

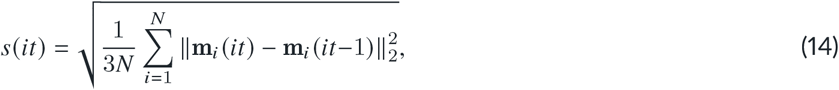

and summarised per run by the mean step size 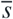 and the final step size *s*_last_. This quantifies how strongly the optimiser had to move the array to achieve its final state.
- **Net drift magnitude:** an overall displacement scale computed from the start and “after” geometries,

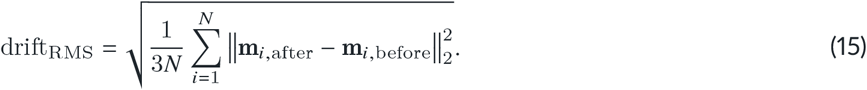

Diagnostics were aggregated by configuration ID using the median across runs (with IQR where appropriate). I additionally visualised *p*_max_, *p*_final_, and *it*_conv_ as grouped boxplots and performed pooled correlation checks between *p*_max_ and the step/drift proxies (to test whether higher attainable pass rates were systematically associated with larger geometry movements). These diagnostics provide an interpretable separation between (i) geometry-dependent limits on achievable localisation performance under the fixed test grid and constraints, and (ii) optimisation dynamics such as convergence speed and stability.

